# Codon bias coevolves with longevity

**DOI:** 10.64898/2025.12.17.694825

**Authors:** Krisztina Kerekes, Mária Trexler, László Bányai, László Patthy

## Abstract

Somatic mutations drive carcinogenesis and aging, shortening the lifespan of animals. Since the vulnerability of genes strongly depends on their size and the abundance of mutation hotspots, we have tested whether negative selection of hypermutable (e.g., CpG bearing) codons could play a role in the evolution of longevity in mammals. Our studies have shown that the CGA codon was significantly more depleted in long-lived than short-lived mammals, suggesting negative selection of this hypermutable stopogenic codon. Interestingly, our analyses have revealed lifespan-dependent changes in codon usage of most amino acids. In the case of a few amino acids (e.g., Ile) the change in codon usage favored translationally optimal codons in long-lived animals, reducing the chances of mistranslation and the formation of abnormal proteins. In the case of a larger group of amino acids (e.g., Tyr, Phe, Asp, Asn), however, the change in codon usage in long-lived animals favored translationally nonoptimal codons that lack matching isodecoder tRNAs. The most likely explanation for this observation is that slowdown of translation at these codons facilitates co-translational folding, thereby reducing the chances of misfolding and aggregation of misfolded proteins in long-lived animals. Our results suggest that the changes in codon usage may contribute significantly to correct co-translational folding, resulting in a more balanced proteostasis and a lower rate of cellular aging in long-lived animals. Our finding is in harmony with the notion that one of the most important hallmarks of aging is loss of proteostasis, manifested in the accumulation of abnormal, misfolded proteins.

## 1. Introduction

Somatic mutations accumulate in protein-coding genes throughout life: they are known to drive carcinogenesis (Stratton et al., 2009) and have long been suspected to play a key role in aging (Szilard, 1959; Morley, 1995). It has been shown recently that in the case of African killifishes relaxed purifying selection and increased mutation load are the prime forces that mold the evolution of the short lifespan of the various species (Cui et al., 2019).

It has been claimed frequently that one major reason why species with extreme longevity avoid cancer and other life shortening diseases is that they have improved repair mechanisms that decrease the rate of somatic mutations that drive carcinogenesis and aging. In a recent study Cagan and coworkers have shown that somatic mutation rates scale with lifespan across mammals. The authors have noted that “the inverse scaling of somatic mutation rates and lifespan is consistent with somatic mutations contributing to ageing” (Cagan et al., 2022). The authors emphasized that „this interpretation is also supported by studies (Zhang et al., 2021) reporting more efficient DNA repair in longer-lived species”.

The role of somatic mutations in carcinogenesis has been clarified in great detail in the last four decades. These studies have shown that mutations of a relatively small fraction of mammalian genes, the „cancer genes” (oncogenes, tumor suppressor genes), play a role in carcinogenesis and only some of the somatic mutations affecting these genes (the „driver mutations”) promote carcinogenesis (Futreal et al., 2004; Sondka et al., 2018). The vast majority of genes are, however, just bystanders: their „passenger” mutations have no role in carcinogenesis (Vogelstein et al., 2013; Bányai et al., 2021). Hanahan and Weinberg have defined several hallmarks of cancer that allow the categorization of cancer genes with respect to their role in the carcinogenesis process (Hanahan and Weinberg, 2011). Cancer genes are usually assigned to hallmarks such as sustained proliferative signaling, evasion of growth suppressors, evasion of cell death, acquisition of replicative immortality, acquisition of capability to induce angiogenesis, activation of invasion and metastasis, tumor promoting inflammation, evasion of immune destruction and reprogramming of energy metabolism. Changes in the maintenance of the genome, epigenome, transcriptome, and proteome occupy a central position in carcinogenesis as they increase the chance that various constituents of cancer hallmarks will acquire capabilities that permit the proliferation, survival, and metastasis of tumor cells (Bányai et al., 2021).

The role of the various mammalian genes in aging is far less clear than in the case of cancer: we do not yet have a widely accepted, annotated census of „aging genes” similar to the Cancer Gene Census (Sondka et al., 2018). Based on the current knowledge about the aging process, López-Otín and coworkers have proposed several aging hallmarks that fulfill the following key criteria: they show age-associated manifestation, their experimental accentuation accelerates aging and therapeutic interventions on these processes decelerate, stop, or reverse aging. Based on these criteria, the authors proposed twelve hallmarks of aging: genomic instability, telomere attrition, epigenetic alterations, loss of proteostasis, disabled macroautophagy, deregulated nutrient-sensing, mitochondrial dysfunction, cellular senescence, stem cell exhaustion, altered intercellular communication, chronic inflammation and dysbiosis (López-Otín et al., 2023a).

It has been recognized a long time ago that carcinogenesis and aging have some common roots in as much as somatic mutations, genomic instability and epigenetic alterations drive both processes. As pointed out by López-Otín and coworkers, aging and cancer are characterized by a series of partially overlapping hallmarks that constitute common “meta-hallmarks” (López-Otín et al., 2023b). The close connection of the two processes may be best illustrated by the fact that DNA repair is a shared hallmark in cancer and ageing. Defects in DNA repair pathways lead to accelerated or premature ageing syndromes and increase the likelihood of cancer development (Clarke and Mostoslavsky, 2022).

Systematic searches for „aging genes” usually employed comparative genomic approaches focusing on long-lived mammalian species such as naked mole rat, bats, whales, elephants (Kim et al., 2011; Seim et al., 2013; Sahm et al., 2018; Tollis et al., 2019; Wilkinson and Adams, 2019; Li et al., 2023; Bowman and Lynch, 2024). The rationale of most of these studies was the assumption that genes/proteins showing strong signs of lifespan-dependent selection are the ones that play key roles in aging. The lists of genes identified in such studies confirmed the close association of longevity and cancer resistance, but they have failed to provide a satisfactory general explanation for the extended lifespan of long-lived species. As pointed out by Brouillette (Brouillette, 2024), the quest to understand aging has long focused on identifying genes responsible for its relentless march. Yet, these efforts have yielded a frustrating lack of genetic clues.”

Recent studies on aging organisms, however, have provided novel insights into the group of genes and types of somatic mutations that may be key etiological factors of aging (for a review see Brouillette, 2024). Soheili-Nezhad and coworkers were the first to find evidence for length-associated transcriptome imbalance in aging organisms. They have shown that in patients with Alzheimer’s disease long genes are more frequently affected by somatic mutations and show reduced expression (Soheili-Nezhad et al., 2021). The authors have pointed out that, if the burden of somatic mutations is uniformly scattered at genomic positions, genes spanning longer portions of the genome are expected to accumulate more mutations than short ones. Accordingly, age-related accumulation of mutations may disproportionally affect longer genes, and this leads to reduced expression of these genes.

Stoeger and coworkers (Stoeger et al., 2022) have shown that the tissues of several vertebrate model organisms display an age-dependent transcriptome imbalance due to significant decrease of transcripts of long genes. They have also noted that in humans and mice the genes with the longest transcripts are enriched in genes reported to extend lifespan, whereas those with the shortest transcripts enrich for genes reported to shorten lifespan. Izeta and coworkers (Ibañez-Solé O, et al., 2023; Soheili-Nezhad at al., 2024), have also demonstrated that in aging organisms there is a negative correlation between gene length and their expression in various cell types, a phenomenon they called length-dependent transcription decline and suggested that this decline contributes to aging as a key etiological factor. They have emphasized that the stochastic nature of somatic mutations provides a plausible explanation why genes are affected proportional to their length. As to the molecular basis of transcription decline the authors suggested that some of the somatic mutations result in transcription-blocking lesions (TBLs) which impede transcriptional elongation and thus have a strong and direct impact on mRNA production (Ibañez-Solé O, et al., 2023).

Length-dependent transcription decline of protein-coding genes in aging organisms, however, is not necessarily due to accumulation of major DNA damages that directly block transcription. Subtle somatic mutations (single nucleotide substitutions, short indels), that interfere with normal functioning of the transcription-translation of genes may also lead to transcription decline. For example, nonsense-mediated mRNA decay that inspects mRNAs for errors that result in the introduction of a premature termination codon, cleaves and eliminates such transcripts from the transcriptome (Kurosaki et al., 2019). Thus, age-dependent decline of transcripts of long genes may be a consequence of the accumulation of nonsense mutations and nonsense mediated decay of their mRNAs. Even nonsynonymous and synonymous single nucleotide substitutions may lead to significant decline of the affected transcripts as codon optimality is a major determinant of mRNA stability (Presnyak et al., 2015). Genome-wide RNA decay analyses have revealed that stable mRNAs are enriched in codons designated optimal, whereas unstable mRNAs contain predominately non-optimal codons. The authors have shown that codon optimality impacts ribosome translocation, connecting the processes of translation elongation and decay through codon optimality.

It was recently shown that the correlation between mRNA and protein levels declines with age in human tissues (Dick et al., 2023; Ding et al., 2025). It has been suggested that progressive uncoupling of transcript and protein levels during cellular ageing may result from increased rates of ribosomal pausing and mistranslation (Llewellyn et al., 2024). In harmony with this interpretation, studies on the effects of aging on the transcriptome and proteome in the brain of short-lived killifish have shown that aberrant translation pausing led to altered abundance of proteins independently of transcriptional regulation (Di Fraia et al., 2025). Interestingly, although aging caused increased ribosome stalling and widespread depletion of proteins in general, stalling events occurred disproportionately on stretches enriched in lysine and arginine codons. The authors point out that the latter observations uncover a potential vulnerable point in the aging brain’s biology: the biogenesis of arginine-rich DNA and RNA binding proteins.

It is important to emphasize that the chances of acquiring nonsense, missense and silent mutations by a protein-coding gene are proportional, not only to its length, but also to the number of hypermutable sites, such as CpG dinucleotides. In our earlier work we have shown that the hypermutability of CpG bearing CGN codons of arginine contribute significantly to the vulnerability of cancer genes (Trexler et al., 2023; Trexler et al., 2024). The CGA codon of arginine is particularly dangerous since, due to hypermutability of its CpG dinucleotide, it generates a nonsense codon (TGA) at an exceptionally high rate (Trexler et al., 2023). These observations raise the possibility that hypermutability of the four CpG bearing codons of arginine might play a significant role in length-dependent transcription decline and progressive uncoupling of transcript and protein levels of aging organisms. It is noteworthy in this respect that in short-lived killifish uncoupling of transcript and protein levels in aging have disproportionately affected arginine-rich DNA and RNA binding proteins (Di Fraia et al., 2025).

In the present work we have tested whether negative selection of hypermutable CpG bearing codons could play a role in the evolution of longevity in mammalian taxa. We have monitored possible signals of selection of synonymous codons by analyzing correlations of codon bias with the lifespan of mammalian species.

Our studies have shown that only the hypermutable CpG bearing stopogenic Arg codon, CGA, was significantly more depleted in long-lived than short-lived mammals, suggesting negative selection of this codon during evolution of longevity. The codon usage of the other CpG-bearing codons, however, did not show significant correlations with longevity of mammals.

Unexpectedly, our analyses have revealed remarkable correlations between evolution of longevity in mammals and changes in usage of synonymous codons of several amino acids. In the case of a few amino acids (e.g., Ile) the change in codon usage favoured translationally optimal codons, i.e. those that match the most abundant isodecoder tRNAs, encoded by the highest number of tRNA genes in mammals. The most likely explanation for this observation is that these translationally optimal codons ensure higher accuracy (lower rate of mistranslation), thereby reducing the chances of the formation of abnormal proteins in long-lived animals (Ke et al., 2017; Bénitière et al., 2025). This explanation is consistent with the fact that one of the most important hallmarks of aging is loss of proteostasis, manifested in the accumulation of abnormal, misfolded proteins (Taylor and Dillin, 2011; Cuanalo-Contreras et al., 2013).

In the case of a larger group of amino acids (e.g. Tyr, Phe, Asp, Asn), however, the change in codon usage favoured translationally nonoptimal codons that lack matching isodecoder tRNA genes in mammals. The most likely explanation for this observation is that slowdown of translation at these codons favours co-translational folding (Sherman and Qian, 2013), thereby reducing the chances of misfolding and aggregation of misfolded proteins in long-lived animals. As pointed out by Pechmann and Frydman, mRNA sequences are under selection to optimize the cotranslational folding of polypeptides (Pechmann S, Frydman J. 2013). According to the hypothesis of these authors nonoptimal codons slow translation to facilitate co-translational folding by allowing the nascent chain more time to develop native-like structure. This explanation is consistent with the fact that one of the most important hallmarks of aging is loss of proteostasis, manifested in the accumulation of misfolded proteins. Our data also provides an explanation for the observation that long proteins play a key role in aging as the chances of misfolding increase with the size and number of folds of multidomain proteins (Rajasekaran and Kaiser, 2022).

In summary, our studies have shown that there are significant lifespan-dependent changes of codon usage that may contribute significantly to improved co-translational folding, resulting in a more balanced proteostasis and a lower rate of cellular aging in long-lived mammals.

## 2. Results and discussion

### 2.1. Impact of GC content of coding genomes on codon bias of mammalian species

To monitor the impact of GC content of coding genomes on codon bias of mammalian species, we have downloaded the RSCU (Relative Synonymous Codon Usage) values of amino acids for placental mammals listed in the Codon Statistics Database (http://codonstatsdb.unr.edu, Subramanian et al., 2022). We limited our study to the 96 mammalian species for which lifespan data are available in the AnAge Database of Animal Aging and Longevity (https://genomics.senescence.info/, de Magalhães et al., 2024).

To determine the overall GC content of the coding sequences (CDSs) for each of the 96 mammalian species, we have downloaded the GC content data of all genes from the Gene Stats section of the Codon Statistics Database (Subramanian et al., 2022). The GC contents of the coding genomes of these species were calculated by weighting the contribution of the genes according to their length (see **Methods**). To visualize the influence of the GC content of coding genomes on codon usage, we have plotted the RSCU values of amino acids as a function of the GC content of the coding genome of the various species, with major emphasis on amino acids Arg, Ser, Thr, Ala, Pro that have at least one codon that contains a CpG dinucleotide. As a control, we have calculated the same parameters for all amino acids, except Trp and Met that are encoded by single codons. We have also calculated the RSCU values of stop codons as a function of the GC content of the coding genome of the various species.

In harmony with earlier conclusions (Plotkin and Kudla, 2011; Trotta, 2016), these analyses have confirmed that the RSCU values of synonymous codons are dictated primarily by the GC content of the coding sequences of the genome. In all cases, the RSCU values of synonymous codons with higher GC content increase, those with lower GC content decrease linearly with GC content of the coding sequences of mammalian species (See **Supplementary file 1** and **Supplementary file 2**). As a typical example we show the results of the analysis of codons encoding alanine (**Figure 1).**

**Figure 1.**
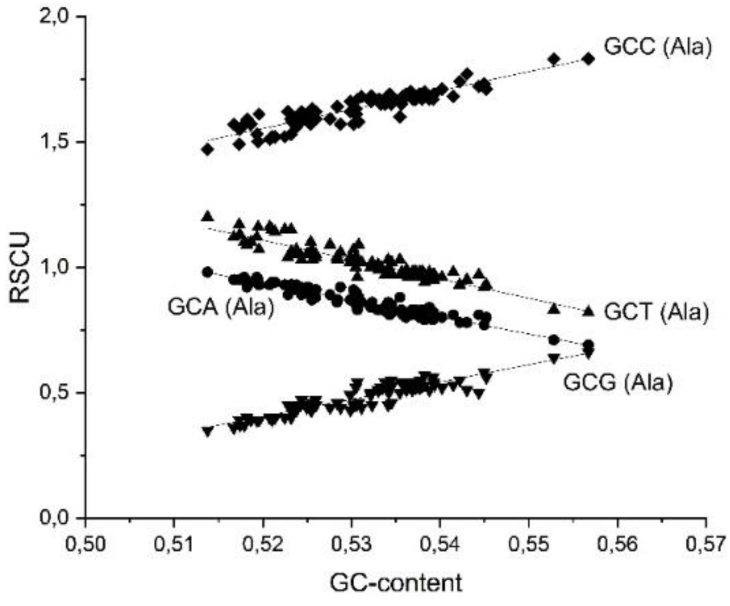
Relative synonymous codon usage (RSCU) of the four alanine codons plotted against the GC content of the coding genomes of the 96 mammalian species included in the present analysis.

In the majority of these linear relationships, the absolute values of Pearson’s r correlation coefficients and R-Square values (COD, Coefficient Of Determination) are very high, close to 1.0 (**Supplementary file 2**), indicating that there is a strong correlation between RSCU values of synonymous codons and the GC content of the coding genome and that the fitted lines explain most of the variability of the response data around its mean.

### 2.2. Impact of hypermutability on codon bias of mammalian species

It is worth pointing out, however, that synonymous codons with similar GC content may have markedly different RSCU values. A major reason for such differences is depletion of codons containing hypermutable sites. This point may be best illustrated by the case of codons containing CpG dinucleotides.

There are eight codons that contain a hypermutable CpG dinucleotide: CGN (Arg) and NCG (Ser, Ala, Thr and Pro). It has been shown earlier that hypermutability of CpG dinucleotides has a significant impact on codon bias of these amino acids in organisms with strongly methylated genomes (Dixon et al., 2016; Trexler et al., 2023; Trexler et al., 2024).

Methylation of cytosines of CpG dinucleotides of the eight codons in the sense strand favors CG>TG mutations resulting in seven missense and one nonsense mutation (Trexler et al., 2024). Methylation of cytosines of CpG dinucleotides of these codons in the antisense strand leads to silent substitutions of the NCG codons of Thr, Pro, Ala and Ser, whereas substitutions of the four CGN codons of Arg lead to missense mutations. In other words, the four Arg codons of the CGN codon family are special among CpG containing codons in that C>T mutation of CpG of both the sense and antisense strand result in a change of the amino acid, whereas in the case of the NCG codons of Thr, Pro, Ala and Ser only the sense strand mutations alter the amino acid. It has been pointed out earlier that this difference of CpG bearing codons has significant evolutionary consequences for organisms with methylated genomes (Dixon et al., 2016). Thanks to purifying selection, NCA codons for Thr, Pro, Ala and Ser increase preferentially with stronger methylation, indicating that depression of NCG codons in strongly methylated genes occurs through silent C>T substitutions on the antisense strand. The CGN group of CpG bearing codons, which code for arginine evolve differently because C>T substitutions on both the sense and antisense stands produce amino acid changes. Dixon and coworkers have shown that arginine content is negatively correlated with gene body methylation, suggesting that the shift in arginine content is due to CpG hypermutability of these codons (Dixon et al., 2016).

As shown in **Supplementary file 2**, the RSCU_50GC_ values (RSCU values calculated for 50% GC content) of CpG bearing codons of Ser, Ala, Pro and Thr are significantly lower than those of their synonymous codons with the same GC-content (Ala: GCC, 1.401 versus GCG, 0.267; Pro: CCC, 1.095 versus CCG, 0.269; Thr: ACC, 1.270 versus ACG, 0.268; Ser: TCC, 1.187 versus TCG, 0.206). In the case of Arg we see much less depletion of CpG bearing codons relative to their GC content equivalent (AGG, 1.354 versus, CGA, 0.734 and CGT, 0.554). These observations are in harmony with the notion that depletion of the CpG bearing NCG codons of Ser, Ala, Pro and Thr occurs by selectively neutral silent substitutions, whereas depletion of CGN codons due to hypermutability of CpG dinucleotides can occur only through nonsense or potentially detrimental missense substitutions.

It is noteworthy, however, that codon usage of CGC and CGG (RSCU_50GC_ values: 0.895 and 0.992, respectively) is less affected by CpG-associated depletion than those of CGA and CGT (RSCU_50GC_ values: 0.734 and 0.554, respectively). The difference in the depletion of these two sets of CGN codons lies in the fact that in the DNA double helix CGC and CGG form three C:G pairs, whereas CGA and CGT form only two C:G pairs, i.e. there is a marked difference in the stability of the double helices formed by these codons. It is highly relevant in this respect that CpG mutation rates produced by 5mC deamination have been shown to be highly dependent on local GC content as DNA melting is the rate-limiting step for 5mC deamination (Fryxell and Moon, 2005). According to this interpretation the higher degree of depletion of CGA and CGT could be due to higher rates of 5mC deamination of these codons. Another factor that might contribute to the more moderate depletion of the GC-rich codons is that mismatch repair is known to be most efficient in regions of stable double helix (Jones et al., 1987).

Hypermutability of the stopogenic CGA codon has another important consequence. As discussed earlier, the fact that out of the codons that can give rise to a stop codon by single nucleotide substitution, CGA is exceptional in as much as it can generate a TGA stop codon at a very high rate (Trexler et al., 2022). This fact explains why in animals with strongly methylated genomes the RSCU value of TGA is much higher than that of its compositional equivalent TAG (Trexler et al., 2022). This is also true for mammals included in the present analysis: the RSCU_50GC_ values of TGA and TAG are 1.416 and 0.677, respectively (**Supplementary file 2**).

### 2.3. Impact of tRNA abundance on codon bias of mammalian species

Synonymous codon usage is known to be under selective pressure to optimize translation efficiency, thus there is a correlation between codon usage and the abundance of cognate tRNAs. Codon bias favouring the most abundant tRNAs is most extreme in highly expressed genes, providing a fitness advantage through increased translation efficiency and accuracy of protein synthesis (Plotkin and Kudla, 2011).

As pointed out by Bénitière and coworkers, tRNA gene copy number is a good predictor of tRNA abundance (Bénitière et al., 2025), therefore we have examined the possible impact of tRNA gene copy number on codon bias (**Supplementary file 2**). Information on the number of tRNA genes in Metazoa, in Mammalia and the group of mammalian species included in the present study have been compiled from data provided by Bénitière and coworkers (Bénitière et al., 2025).

In Metazoa the 18 amino acids that are encoded by multiple synonymous codons form two major groups: (i) those whose synonymous codons are translated by more than one distinct isodecoder tRNAs; (ii) those for which all synonymous codons are translated by a single isodecoder tRNA (Bénitière et al., 2025). The first group includes amino acids encoded by codon sextets (Leu, Arg and Ser), quartets (Val, Gly, Ala, Pro and Thr), a triplet (Ile) and NNG/NNA duets (Glu, Gln and Lys). NNG/NNA duets are unique in that each synonymous codon has a matching isodecoder tRNA, whereas in the case of amino acids with codon sextets, quartets and triplet at least one of the codons has no matching isodecoder tRNA.

The second group for which all synonymous codons are translated by a single isodecoder tRNA corresponds to the six amino acids encoded by NNC/NNT duets (Phe, Cys, Tyr, Asp, His, Asn).

Differences in the number of the various isodecoder tRNA genes (abundance or complete absence of tRNA species) may have a major impact on codon bias. This point may be best illustrated by the case of the CTC and CTG codons of Leucine (**Supplementary file 2**). Despite the similarity of their GC content, the RSCU values of CTC and CTG, extrapolated to GC content of 0.5, are markedly different: 1.084 and 2.100, respectively. It seems most likely that the strong preference of CTG over CTC is explained by the fact that the median copy number of isodecoder tRNA genes for CTG is the highest in most mammalian species (**Supplementary file 2**), whereas isodecoder tRNA genes for CTC are missing (Bénitière et al., 2025).

### 2.4. Changes in codon usage and evolution of longevity in rodent species

To test the assumption that negative selection of hypermutable CpG bearing codons could play a role in the evolution of longevity in mammals, we have first focused on changes of codon usage of the codon families CGN (Arg) and NCG (Ser, Ala, Thr and Pro) as a function of the lifespan of mammalian species, but we have extended these studies to all other amino acids.

For each species, we have subtracted the RSCU values expected at the given GC content (RSCU_exp_) from the actual RSCU values (RSCU_obs_) and plotted these deviations as a function of the known lifespan (years) of the species (**Supplementary file 3**).

To minimize the influence of major taxon-specific biological differences on such a hypothetical correlation, we have first restricted our analyses to 19 rodent species then we have checked whether the correlations observed in the case of rodents are also detectable across a wider range of mammals. The justification for this approach is that some taxon-specific biological differences are known to have greater influence on mortality and lifespan than cellular aging. For example, bats have several unique features (powered flight, viral tolerance and disease resistance, hibernation. torpor) that contribute to longer lifespan explaining why most bats have lifespans more than fourfold of similar-sized placental mammals (Wilkinson and Adams, 2019; Morales et al., 2025).

In rodents the majority of amino acids had significant lifespan dependent changes of codon usage. There were strong (absolute Pearson’s r values exceeding 0.5) or moderate (absolute Pearson’s r values exceeding 0.3) correlations between lifespan and change in codon usage in the case of most amino acids (**Supplementary file 2**).

In the case of CpG-bearing codons of the NCG group, there was a significant linear correlation between lifespan and depletion of CCG (Pearson’s r = –0.4436), TCG (Pearson’s r= – 0.5724) and ACG (Pearson’s r=-0.6549), suggesting that decrease in the number of these mutation hotspots favours longevity of rodents.

In the case of the CGN codons, there was a significant linear correlation between lifespan and depletion of CGA (Pearson’s r =-0.6096) and CGT (Pearson’s r=-0.6007) and increased use of CGC (Pearson’s r = 0.6438) and CGG (Pearson’s r=0.6151). A possible explanation for these observations is that depletion of the CGA and CGT codons occurs primarily by silent CGA, CGT>CGC, CGG substitutions and that substitution of the CGA, CGT codons with the less vulnerable synonymous CGC, CGG codons (see section 2.2.) favours longevity of rodents.

Significant changes of codon usage were also observed in the case of several NNC/NNT duets that have a single isodecoder tRNA.

Our analyses have revealed significant correlations between lifespan of rodents and changes of codon preference of Asn (Pearson r = ±0.7867), Asp (Pearson r = ±0.6639), Cys (Pearson r =±0.5259), Phe (Pearson r =±0.7530) and Tyr (Pearson r = ±0.7446). As a typical example, we show the results of the analyses of the data for the two codons of phenylalanine (**Figure 2, panels A and B).**

**Figure 2.**
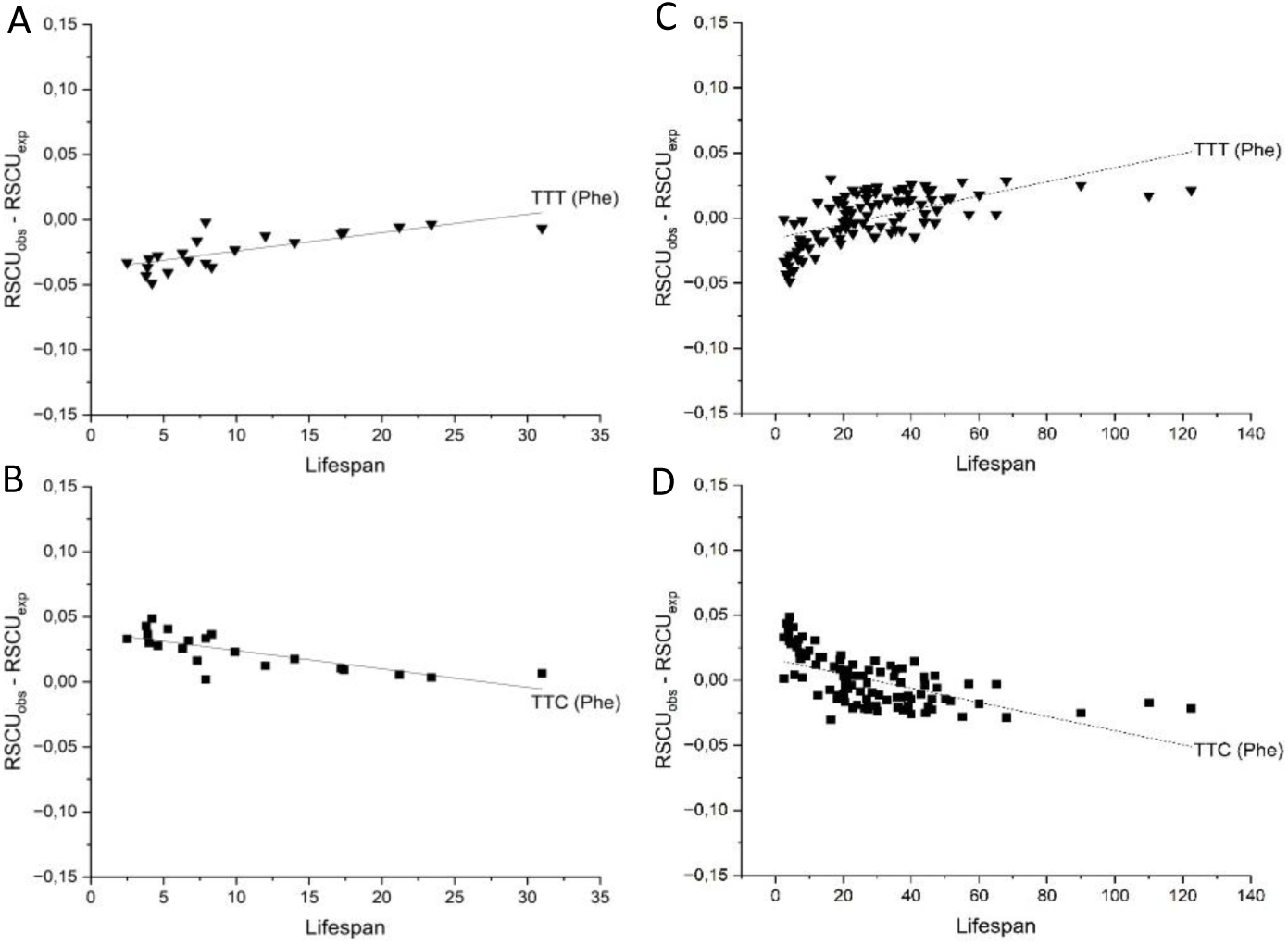
Correlation between lifespan of mammalian species and changes in codon preference of phenylalanine (Phe). Panels A and B show data for the 19 rodent species, Panels C and D show data for the 96 mammalian species included in the present analysis.

Interestingly, in the case of Asn, Asp, Cys, Phe and Tyr the change in codon usage favoured the use of the NNT codons that have no matching isodecoder tRNA gene in mammals and disfavoured the translationally optimal NNC codons that have matching isodecoder tRNA genes (**Supplementary file 2**). The most likely explanation for this observation is that slowdown of translation at some nonoptimal Asn, Asp, Cys, Phe, Tyr codons favours co-translational folding (Sherman and Qian, 2013; Pechmann and Frydman J, 2013), thereby reducing the chances of misfolding and aggregation of misfolded proteins in long-lived rodents.

Change of codon usage favouring translationally nonoptimal codons over optimal codons is not restricted to the set of NNC/NNT duets. In the case of Ala, it is the GCT codon that has the highest number of cognate tRNA genes, whereas the GCC codon lacks matching isodecoder tRNA genes. We have observed a significant correlation between lifespan and reduced use of the optimal GCT codon (Pearson’s r = –0.6105), whereas the use of the nonoptimal GCC codon showed a weak lifespan-dependent increase (Pearson’s r = 0.2447).

In the case of Arg, it is the CGC codon that lacks cognate tRNA genes, whereas the CGT codon is the one that has the highest number of cognate tRNA genes. We have observed a significant correlation between lifespan and reduced use of the optimal CGT codon (Pearson’s r = –0.6007), whereas the use of the nonoptimal CGC codon showed a lifespan-dependent increase (Pearson’s r = 0.6438). It seems possible that reduced use of CGT in long-lived rodents has more to do with optimalization of co-translational folding than with elimination of CGT as a mutation hotspot.

In the case of Pro, it is the CCC codon that lacks isodecoder tRNA genes, whereas CCT has the highest number of tRNA genes. We have observed a significant correlation between lifespan and reduced use of the optimal CCT codon (Pearson’s r = –0.3885), whereas the use of the nonoptimal CCC codon showed a significant lifespan-dependent increase (Pearson’s r = 0.4932).

AGT is one of the serine codons that lacks isodecoder tRNA genes, whereas the TCT codon of serine has the highest number of tRNA genes. We have observed a significant correlation between lifespan of rodents and reduced use of the optimal TCT codon (Pearson’s r = –0.6309), whereas the use of the nonoptimal AGT codon showed significant lifespan-dependent increase (Pearson’s r = 0.5532).

Several other amino acids, however, showed an opposite phenomenon: the change of codon usage favoured the translationally optimal codons that have the highest number of matching isodecoder tRNA genes and disfavoured the use of codons that have no matching isodecoder tRNA gene in mammals (**Supplementary file 2**). Changes of codon usage of Gly may illustrate this point. The Gly codon, GGC has the highest number of tRNA genes, whereas the GGT codon lacks cognate isodecoder tRNA genes. In this case, we have observed a significant correlation between lifespan of rodents and reduced use of the nonoptimal GGT codon (Pearson’s r = –0.4979), whereas the use of the optimal GGC codon showed a lifespan-dependent increase (Pearson’s r = 0.4576).

This phenomenon is most striking in the case of Ile (**Figure 3**). Out of the three codons of this amino acid the ATT codon has the highest number of matching isodecoder tRNA genes, whereas the ATC codon has no matching isodecoder tRNA gene in mammals (**Supplementary file 2**). We have observed a significant correlation between lifespan of rodents (**Figure 3. Panels A and B**).and reduced use of the nonoptimal ATC codon (Pearson’s r = –0.7885), whereas the use of the optimal ATT codon showed a lifespan-dependent increase (Pearson’s r = 0.7590).

**Figure 3.**
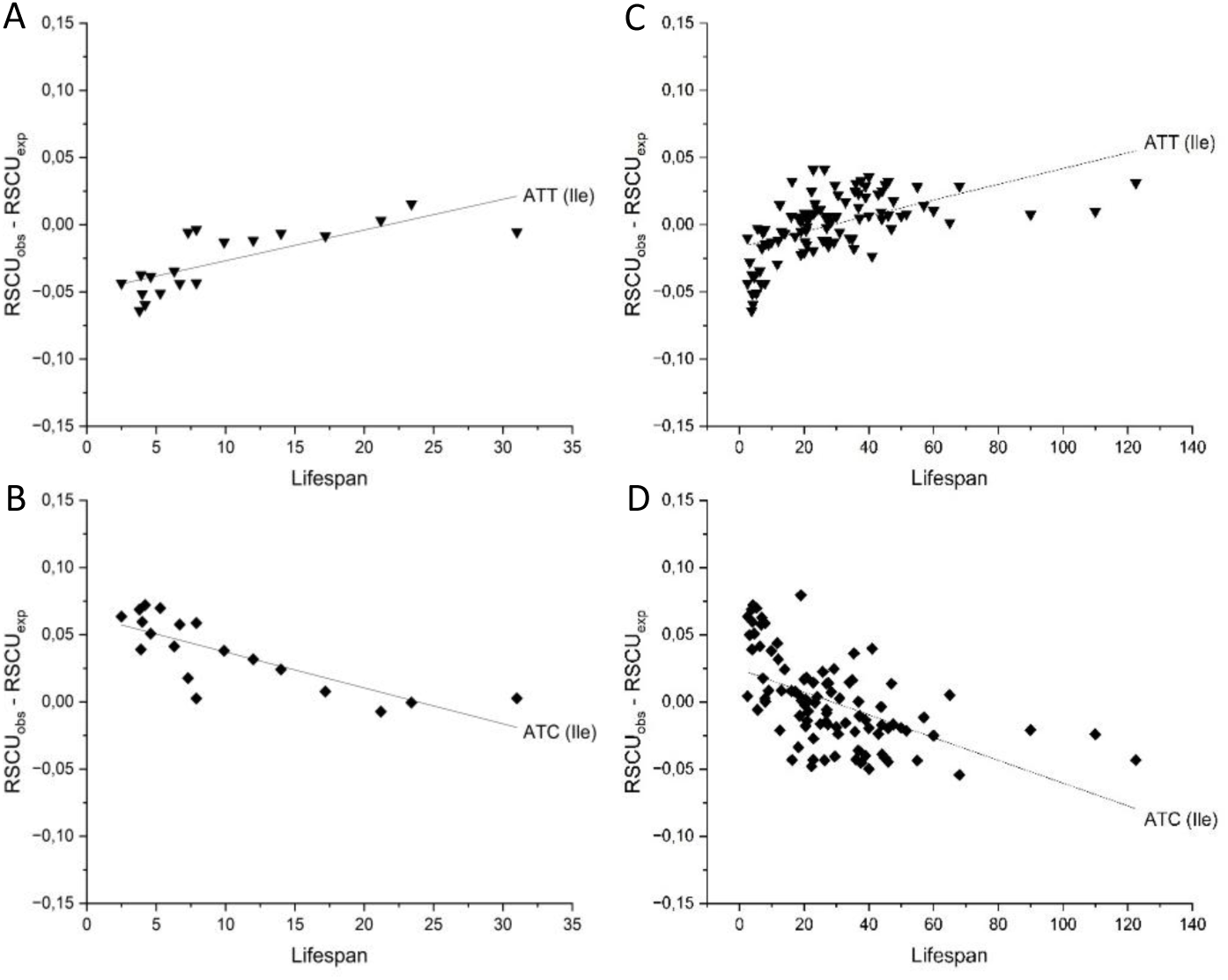
Correlation between changes in codon preference of isoleucine (Ile) and lifespan of mammalian species. Panels A and B show data for the 19 rodent species, Panels C and D show data for the 96 mammalian species included in the present analysis.

In the case of Leu, the CTG codon has the highest number of matching isodecoder tRNA genes in mammals, whereas the CTC codon that has no matching isodecoder tRNA gene. We have observed a significant correlation between lifespan of rodents and reduced use of the nonoptimal CTC codon (Pearson’s r =-0.7331), whereas the use of the optimal CTG codon showed a lifespan-dependent increase (Pearson’s r = 0.5497).

Changes of codon usage of Thr also suggest that evolution of longevity of rodents may favour selection for translationally optimal codons. In the case of this amino acid, ACT codon has the highest, ACG has the lowest number of tRNA genes. We have observed a significant correlation between lifespan and reduced use of the less optimal ACG codon (Pearson’s r = – 0.6549), whereas the use of the optimal ACT codon showed a lifespan-dependent increase (Pearson’s r = 0.6771).

Evolution of longevity of rodents may also favour selection for translationally optimal Val codons. In the case of Val, the GTG codon has the highest number of matching isodecoder tRNA genes, whereas the GTC codon has no matching isodecoder tRNA gene in mammals. We have observed a significant correlation between lifespan and reduced use of the nonoptimal GTC codon (Pearson’s r = –0.7723), whereas the use of the optimal GTG codon showed a lifespan-dependent increase (Pearson’s r = 0.5407).

The NNG/NNA duets (Glu, Gln, Lys), where both synonymous codons have a comparable number of matching isodecoder tRNA genes do not fit into either of the two categories listed above: there is no clear distinction between optimal and nonoptimal codons. We observed no significant correlation between change of codon usage and lifespan of Gln. Long-lived rodents, however, slightly favoured the use of GAA codon of Glu over GAG codon (Pearson’s r = ±0.3372). Since the number of isodecoder tRNA genes for GAG and GAA are similar, the slight shift of codon preference is unlikely to be explained by differences in their optimality for translation or cotranslational folding. Since GAG triplet repeat sequences may form hairpin structures and may thus cause triplet repeat expansion diseases (Mitas, 1997) it seems possible that depletion of GAG may reflect negative selection in long-lived rodents.

Surprisingly, in the case of Lys, there was a strong correlation between changes in codon preference and lifespan of rodents (Pearson’s r =±0,736). Since in most mammals the numbers of genes of isodecoder tRNA for AAG and AAA are similar the change of codon preference favouring the AAA codon over AAG, is unlikely to be explained by differences in their optimality for translation and folding of proteins. It is more likely that this change in codon preference is a reflection of the strong negative selection of AAG trinucleotide repeats in exons (Kozlowski et al., 2010).

In summary, our analyses have identified three distinct groups of amino acids that show lifespan-dependent changes of codon bias in rodents: 1) amino acids, where there is a correlation between increased use of nonoptimal codons and increased lifespan of rodents (e.g. Asn, Asp, Cys, Phe, Tyr, Ala, Arg, Pro, Ser); 2) amino acids, where there is a correlation between increased use of optimal codons and increased lifespan of rodents (e.g. Gly, Ile, Leu, Thr, Val); 3) amino acids, where there’s no clear distinction of optimal and nonoptimal codons (Glu, Lys).

The most likely explanation for the increased use of translationally optimal codons in long-lived rodents is that this change ensures increased accuracy of translation (Sun and Zhang, 2022) and the lower rate of mistranslation results in a more balanced proteostasis and a lower rate of cellular aging in long-lived rodents. This interpretation is in harmony with recent studies on rodent species with diverse lifespans that have shown that the fidelity of protein synthesis is a determining factor in the evolution of longevity (Azpurua et al., 2013; Ke et al., 2017; Ke et al., 2018); the basal rate of translation errors have shown strong negative correlation with maximum lifespan.

Increased use of translationally nonoptimal codons of Asn, Asp, Cys, Phe, Tyr, Ala, Arg, Pro, Ser in long-lived rodents was more surprising as nonoptimal codons may increase the chance of mistranslation, ribosome stalling and this can result in premature translation termination (Ke et al., 2017). Use of nonoptimal codons, however, may have significant benefits if slowdown of translation at such codons, favours correct co-translational folding by allowing the nascent chain more time to develop native-like structure before continuing translation (Pechmann and Frydman, 2013). This hypothesis implicitly assumes that such a beneficial role of nonoptimal codons of protein-coding genes is „position-specific”: they work best if they occur in regions separating distinct folding units. It is noteworthy in this respect that Pechmann and Frydman have noted that hydrogen-bonded turns that frequently connect secondary structural elements (α-helices and β-strands) are strongly enriched in conserved nonoptimal codons (Pechmann and Frydman, 2013). This principle is also valid for linker regions that connect domains of multidomain proteins. As pointed out by Jacobson and Clark, positioning of nonoptimal codons in linker regions could facilitate a domain-by-domain, beads-on-a-string folding mechanism (Jacobson and Clark, 2016).

We suggest that correct co-translational folding of multidomain proteins in long-lived rodents is facilitated by increased use of nonoptimal codons in the linker regions that connect the individual domains of these proteins. Since multidomain proteins constitute a very large fraction of the proteomes of animals (Tordai et al., 2005) and co-translational folding of multidomain proteins is particularly challenging (Rajasekaran and Kaiser, 2022) we suggest that the increased use of nonoptimal codons in these regions contributes significantly to longevity of rodents.

### 2.5. Changes in codon usage and evolution of longevity in mammalian species

In the next part of our work, we have examined whether the various lifespan-dependent changes in codon usage observed in the analysis of rodents are detectable when we analyse a broader spectrum of mammals with greater biological diversity.

Comparison of the correlations between lifespan and changes in codon usage in the analysis of mammals with those observed when the analysis was limited to rodents revealed that several correlations significant in rodents are obscured when we analyse a more diverse group of mammals. Several correlations remained significant but in the majority of cases Pearson’s r values of the correlations were lower in the analysis of mammals (**Supplementary file 2).**

The correlations between mammalian lifespan and changes in codon usage deviate most significantly from linearity in the case of mammalian species with extreme longevity. As shown in panels C and D of **Figures 2** and **3**, the slopes of the linear changes in codon usage are the steepest in the range of relatively short-lived mammals (lifespan range of 1-40 years), less steep in the range of moderately long-lived mammals (lifespan range of 41-60 years) and practically zero in the case of extremely long-lived mammals (lifespan range above 60 years). The hyperbolic curves reflecting the correlation of changes in codon usage and lifespan suggest that although the change of codon usage contributes to the evolution of extended lifespan of mammals, extreme longevity of species such as humans and whales are not due to even more extreme shifts in codon usage. The most likely explanation for this phenomenon is that during evolution of increased lifespan changes in codon usage saturate the sites critical for efficient translation or folding of multidomain proteins, reaching a point in long-lived mammals where lifespan becomes limited by processes other than protein misfolding and collapse of proteostasis.

Whereas in rodents there was a significant correlation between lifespan and depletion of the CpG bearing codons CCG, TCG and ACG, no such depletion was evident from the analysis of a broader spectrum of mammals. This observation suggests that depletion of these mutation hotspots does not play a general role in the longevity of mammals. The significant correlation between lifespan and depletion of the arginine codon, CGA observed in rodents was, however, also observed in mammals (Pearson’s r = –0.4051) concomitant with the increased use of CGG (Pearson’s r=0.3681). These observations suggest that depletion of the hypermutable stopogenic CGA codon occurs primarily by silent CGA>CGG substitution and that substitution of the stopogenic CGA codon with a synonymous nonstopogenic codon favours longevity of mammals.

Similarly to rodents, significant changes of codon usage were observed in the case of several NNC/NNT duets that have a single isodecoder tRNA. Our analyses have revealed a linear relationship between change of codon preference and lifespan in the case of Asn (Pearson r = ±0.4419), Asp (Pearson r = ±0.5123), Phe (Pearson r = ±0.5994) and Tyr (Pearson r = ±0.4747). In all these cases the change in codon usage favoured the use of the nonoptimal NNT codon that has no matching isodecoder tRNA gene in mammals (**Supplementary file 2**). In the case of the NNC/NNT duets, Cys and His, however, we observed only very weak correlation between change of codon usage and lifespan (**Figure 2, panels C and D**). We have observed a weak but significant correlation between lifespan of mammals and reduced use of the optimal GCT codon of Ala (Pearson’s r = –0.3779), whereas the use of the nonoptimal GCC codon showed a weak lifespan-dependent increase (Pearson’s r = 0.2187). We have observed a significant correlation between lifespan of mammals and reduced use of the optimal CCT codon of Pro (Pearson’s r = –0.4139), whereas the use of the nonoptimal CCC codon showed a lifespan-dependent increase (Pearson’s r = 0.2087). In contrast with the rodents-analysis, we have observed no significant correlation between mammalian lifespan and reduced use of optimal and increased use of nonoptimal codons of Ser and Arg.

Change of codon usage favouring the translationally optimal codons (and disfavouring the use of codons that have no matching isodecoder tRNA gene) was far more restricted across mammals than in the case of rodents-analysis. We have observed no significant correlation between lifespan of mammals and reduced use of the nonoptimal and increased use of the optimal codons of Gly, Leu, Thr and Val. Only in the case of Ile did we observe a significant correlation between mammalian lifespan and reduced use of the nonoptimal ATC (Pearson r = – 0.5485 and increased use of the optimal the ATT codon (Pearson r = 0.5407) (**Figure 3, panels C and D**).

The results on the NNG/NNA duets (Glu, Gln, Lys) where each synonymous codon has a comparable number of matching isodecoder tRNA genes, were quite similar in the two analyses. In the case Gln we observed a weak correlation between change of codon usage and lifespan (Pearson r = 0.2191), long-lived mammals preferring the CAA over CAG. Long-lived mammals also favour the use of GAA codon of Glu over GAG codon (Pearson’s r = 0.3976). Analyses of mammals also revealed a strong correlation between changes in codon preference of Lys and lifespan (Pearson’s r = 0.5412).

In summary, our analyses have shown that several of the significant lifespan-dependent changes in codon usage observed in rodents are also observed when we analyse a broader spectrum of mammals. Significantly, there is a strong correlation between increased use of nonoptimal codons and increased lifespan of mammals in the case of Asn, Asp, Phe, Tyr, Ala, and Pro whereas there is a strong correlation between increased use of optimal codon and mammalian lifespan only in the case of Ile. Since preference of translationally nonoptimal codons may be essential for correct co-translational folding (Jacobson and Clark, 2016) this observation suggests that correct protein folding is more critical for mammalian longevity than high rate of translation that would favour translationally optimal codons.

Since multidomain proteins constitute a very large fraction of the proteomes of animals and much of their organismic complexity is ensured by these proteins (Tordai et al., 2005) their correct folding is of extreme importance. The folding of multidomain proteins, however, differs from that of small single-domain proteins, which usually fold quickly and reversibly. Co-translational processes are important aspects of multi-domain protein folding: as the ribosome sequentially adds amino acids from N-to C-terminus, the growing nascent polypeptide begins to form tertiary structures, domain by domain (Jacobson and Clark, 2016; Rajasekaran and Kaiser, 2022; Rajasekaran and Kaiser, 2024). We suggest that increased use of translationally nonoptimal codons in linker regions of multidomain proteins favours their correct co-translational folding and may thus play a critical role in the evolution of mammalian longevity.

Although our work focused on codon bias of synonymous sense codons, we note with interest that there is a significant correlation between mammalian lifespan and increased use of the TAA stop codon (Pearson r = 0.4333) and decreased use of TAG (Pearson r = –0.4279) and TGA (Pearson r = –0.2542) as stop codons (**Supplementary file 2**). It is now well established that the three stop codons are not fully synonymous as they differ in efficiency to terminate translation (Trexler et al., 2023). TAA appears to be the most efficient, whereas the others permit significant readthrough due to misinterpretation of the stop codon. As we have discussed earlier (Trexler et al., 2023), it has been debated for a long time whether readthrough of termination codons is detrimental (as it creates aberrant forms of proteins) or has adaptive value (as it provides a means for proteome expansion). Our observation that evolution of mammalian longevity favors the use of TAA that is the least prone to readthrough suggests that avoidance of the formation of aberrant forms of proteins is of primary importance for lower rates of cellular aging.

## 3. Conclusion

In the present work we have detected significant correlations between evolution of lifespan in mammals and changes in codon usage of the majority of synonymous codons.

For example, the hypermutable CpG bearing stopogenic arginine codon, CGA, is significantly more depleted in long-lived than short-lived mammals. The most likely explanation for this observation is that negative selection of this codon renders genes less vulnerable (Trexler et al., 2024; Di Fraia et al., 2025), promoting the evolution of longevity.

We have also noted that there is a significant correlation between extended mammalian lifespan and increased use of the TAA stop codon (and decreased use of TAG and TGA as stop codons). Since TAA is the least prone to stop codon readthrough, this observation suggests that avoidance of the formation of abnormal proteins with aberrant C-terminal extensions is important for lower rates of cellular aging.

In addition to these special cases, our analyses have revealed remarkable correlations between evolution of longevity in mammals and changes in usage of the majority of synonymous codons.

In the case of some amino acids (e.g., Ile) the change in codon usage accompanying evolution of longevity favoured translationally optimal codons, i.e. those that match the most abundant isodecoder tRNAs. The most plausible explanation for this observation is that these translationally optimal codons ensure higher accuracy (lower rate of mistranslation), thereby reducing the chances of the formation of abnormal proteins in long-lived animals (Ke et al., 2017; Bénitière et al., 2025).

Unexpectedly, in the case of a much larger group of amino acids (e.g., Tyr, Phe, Asp, Asn), the change in codon usage accompanying evolution of longevity favoured translationally nonoptimal codons that lack matching isodecoder tRNA genes. In our view, the most likely explanation for this observation is that slowdown of translation at these codons favours co-translational folding as suggested earlier (Sherman and Qian, 2013; Pechmann and Frydman, 2013; Jacobson and Clark, 2016). According to the hypothesis of these authors, nonoptimal codons slow translation and facilitate co-translational folding by allowing the nascent chain more time to develop native-like structure.

Co-translational processes are especially important for folding of multi-domain proteins as the growing polypeptide must form tertiary structures domain by domain (Jacobson and Clark, 2016; Rajasekaran and Kaiser, 2022; Rajasekaran and Kaiser, 2024). We suggest that increased use of translationally nonoptimal codons in critical regions of proteins (such as linker regions of multidomain proteins) favours their correct co-translational folding and may thus play a critical role in the evolution of mammalian longevity (**Figure 4**). This explanation is consistent with the fact that one of the most important hallmarks of aging is loss of proteostasis, manifested in the accumulation of misfolded proteins.

**Figure 4.**
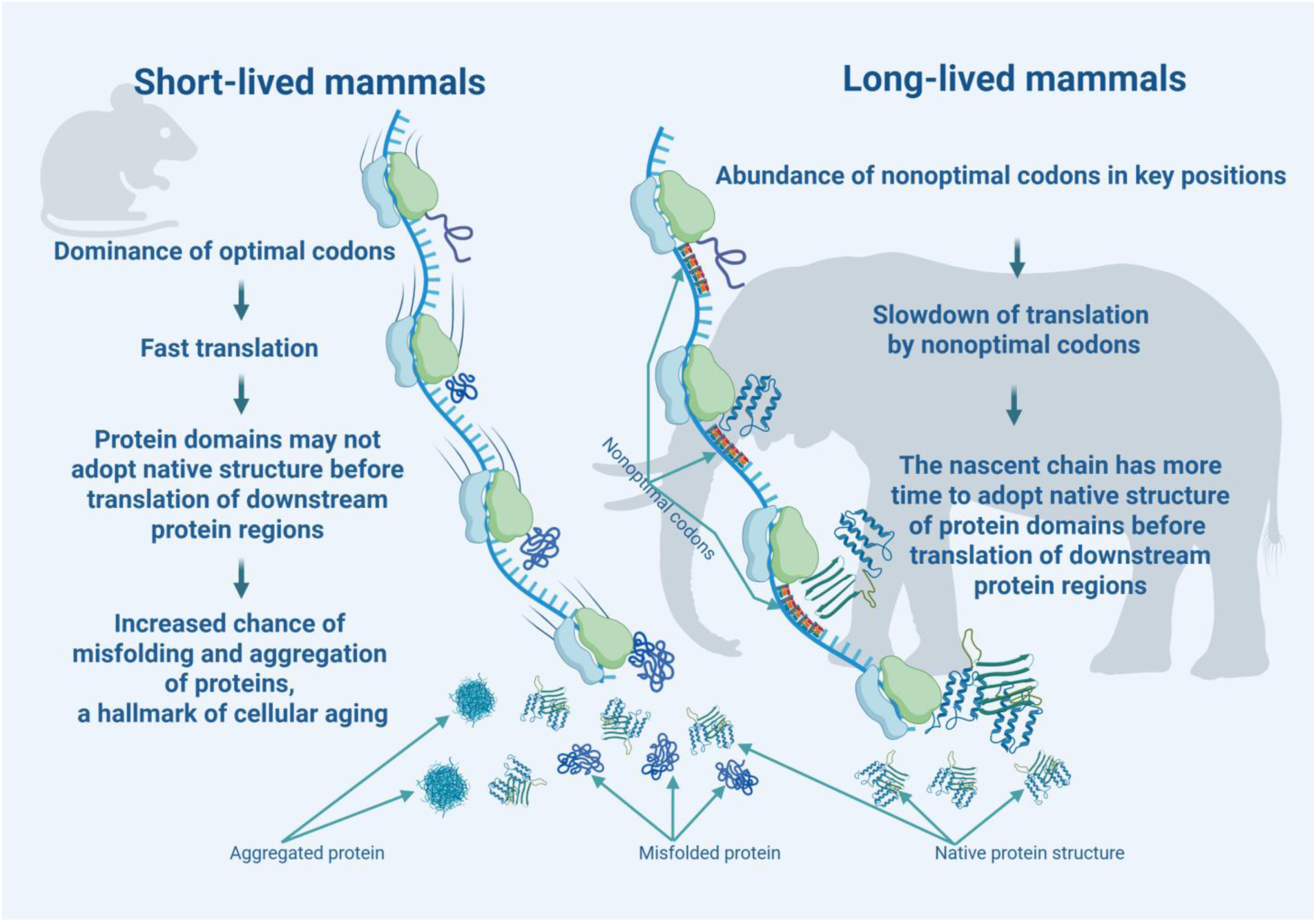
Proposed model for the effects of optimal and nonoptimal codons on folding of proteins and longevity. Created with BioRender.com. In short-lived mammals translation is fast due to the dominance of optimal codons that match the most abundant isodecoder tRNAs, therefore the protein domains may not adopt their stable native structure before translation of downstream protein regions. High speed of translation may increase the chance of misfolding and aggregation of misfolded proteins, a hallmark of cellular aging. In long-lived mammals nonoptimal codons replace optimal codons in critical regions (e.g., linker regions of multidomain proteins). Slowdown of translation at these codons favours co-translational folding by allowing the nascent chain more time to develop native-like structure before translation of downstream protein regions. Thus, replacement of optimal codons by nonoptimal codons in critical regions promotes co-translational folding and decreases the chance of misfolding and aggregation of misfolded proteins.

## 4. Methods

### 4.1. Lifespan data of mammalian species

Lifespan information (maximum lifespan values) for mammalian species was retrieved from the AnAge Database of Animal Aging and Longevity (https://genomics.senescence.info/, de Magalhães et al., 2024)

### 4.2. Codon usage and genomic GC content data of mammalian species

The relative synonymous codon usage (RSCU) values for the coding sequences of mammalian species were downloaded from the Codon Statistics Database (http://codonstatsdb.unr.edu, Subramanian et al., 2022).) over the period December 3–13, 2024. We limited our study to the 96 mammalian species for which complete datasets were available in both the Codon Statistics Database and the AnAge Database.

To determine the overall GC content of the coding genome for the 96 mammalian species, we have downloaded the GC content data of genes from the Gene Stats section of the Codon Statistics Database (**Supplementary file 4**). The GC contents of the coding genomes of these species were calculated by weighting the contribution of the genes according to their length. For each gene of a given species the GC fraction values were multiplied by the number of codons present in the gene and the sum of these values of all genes were divided by the total number of codons of all genes.

To visualize the influence of the GC content of coding genomes on codon usage, we have plotted the RSCU values of amino acids as a function of the GC content of the Coding Genome of the various species.

### 4.3. Data analysis and statistical procedures

To monitor the possible correlations between changes in codon bias and lifespan we have calculated the RSCU values expected at the given GC content (RSCU_exp_), based on the linear relationship between the RSCU and GC content. RSCU_exp_ values were subtracted from the actual RSCU values (RSCUobs), and we have plotted these deviations as a function of the known lifespan (years) of the species.

All datasets were processed and analyzed using OriginPro 2025. Regression analyses and Pearson correlation computations were performed to assess the relationships between codon usage metrics, GC content and species lifespan. In the evaluation of the results of these analyses we have focused on strong correlations, i.e. cases where the absolute value of Pearson correlation coefficient (r) was higher than 0.5. Cases where the absolute value of Pearson correlation coefficient (r) was between 0.3 and 0.5 are discussed as moderate but significant correlations.

## Supporting information

**Supplementary file 1.** Impact of GC content of coding genomes on codon bias of mammalian species

**Supplementary file 2.** Copy number of tRNA genes in metazoa and impact of GC content of coding genomes and lifespan on relative synonymous codon usage (RSCU) of mammalian species

**Supplementary file 3.** Changes in codon usage and evolution of longevity in mammalian species

**Supplementary file 4.** GC content of the coding genomes and relative synonymous codon usage (RSCU) of 96 mammalian species with known lifespan

## Acknowledgments

KK, MT, LB and LP are supported by the GINOP-2.3.2-15-2016-00001 grant of the Hungarian National Research, Development and Innovation Office (NKFIH). We appreciate the tools of BioRender that were instrumental for the visualization of our findings.

## Contributor Information

Krisztina Kerekes

Mária Trexler

László Bányai

László Patthy

Krisztina Kerekes, Mária Trexler, László Bányai: Formal analysis, Validation, Investigation, Methodology, Writing—original draft, Writing—review and editing; László Patthy, Conceptualization, Supervision, Funding acquisition, Validation, Methodology, Writing—original draft, Project administration, Writing—review and editing.

## Data availability

All data generated or analysed during this study are included in the manuscript and supporting files.

