## Supplementary file 1 for "Codon bias coevolves with longevity"

### Impact of GC content of coding genomes on codon bias of mammalian species

The file presents the plots reflecting the impact of GC content of coding genomes on codon bias of mammalian species. The parameters of the regression analyses of these plots are presented in Supplementary file 2.

The plots are presented in alphabetical order of the names of the amino acids; data for amino acids Met and Trp that are encoded by a single codon are not shown.

We also present the data for the three stop codons at the end of the list of the 18 amino acids encoded by more than one synonymous codon.

In the case of the 18 amino acids and the termination codons the figures present the plots reflecting the impact of the GC content of the coding genome of mammalian species on the relative synonymous codon usage (RSCU) of the given amino acid or termination codons. In these analyses we have plotted the RSCU values as a function of the GC-content of the coding genomes of the 96 mammalian species included in the present analysis.

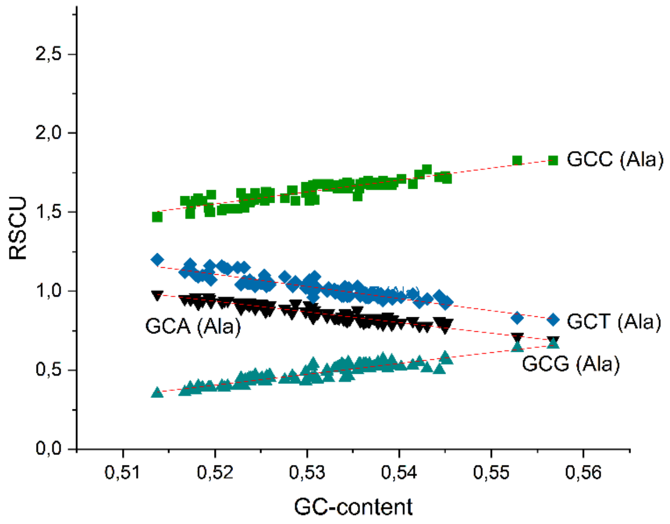


**Supplementary figure 1.a.** Impact of the GC content of coding genomes of mammalian species on their relative synonymous codon usage (RSCU) of the four **alanine** codons.

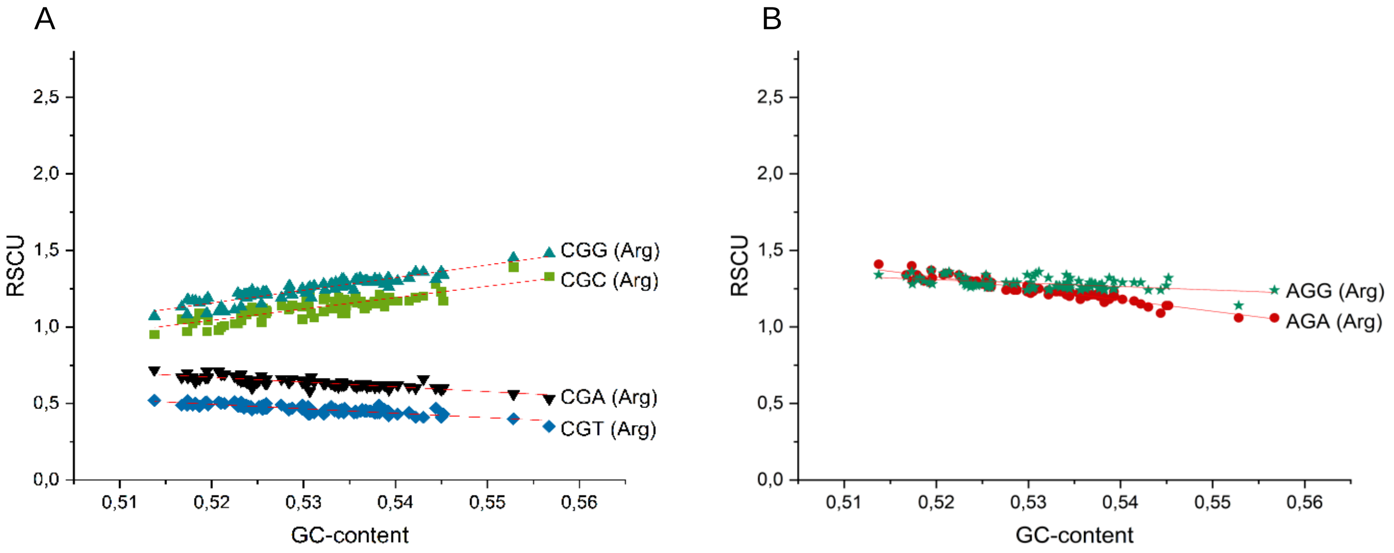


**Supplementary figure 2.a.** Impact of the GC content of coding genomes of mammalian species on their relative synonymous codon usage (RSCU) of the six **arginine** codons.

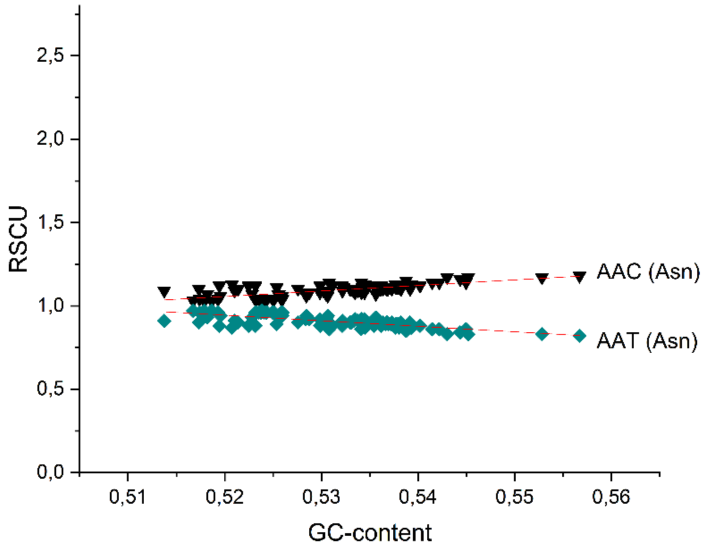


**Supplementary figure 3.a.** Impact of the GC content of coding genomes of mammalian species on their relative synonymous codon usage (RSCU) of the two **asparagine** codons.

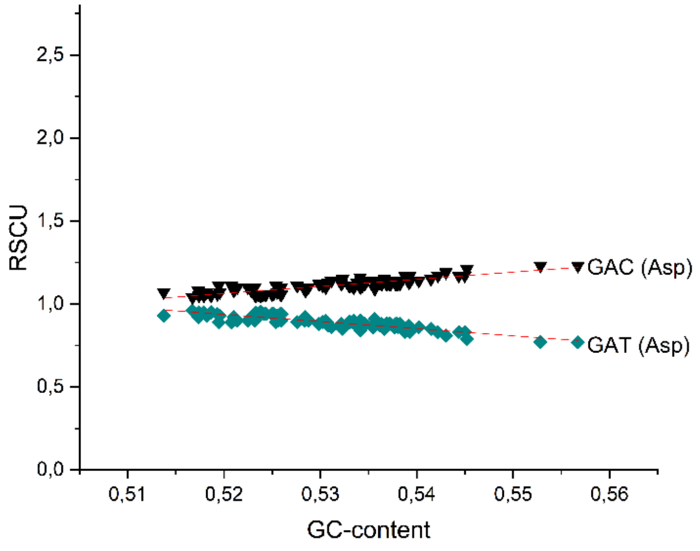


**Supplementary figure 4.a.** Impact of the GC content of coding genomes of mammalian species on their relative synonymous codon usage (RSCU) of the two **aspartic acid** codons.

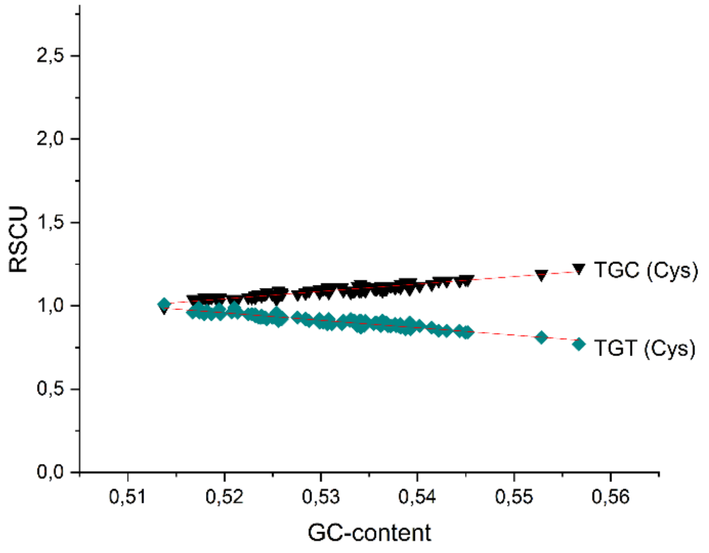


**Supplementary figure 5.a.** Impact of the GC content of coding genomes of mammalian species on their relative synonymous codon usage (RSCU) of the two **cysteine** codons.

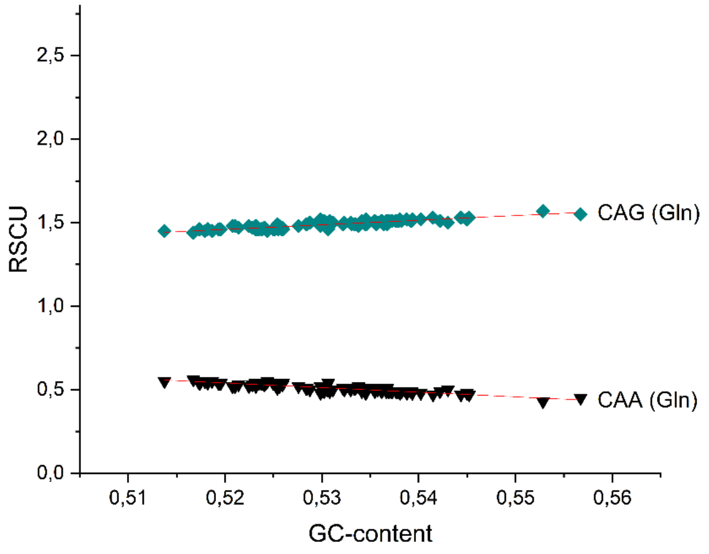


**Supplementary figure 6.a.** Impact of the GC content of coding genomes of mammalian species on their relative synonymous codon usage (RSCU) of the two **glutamine** codons.

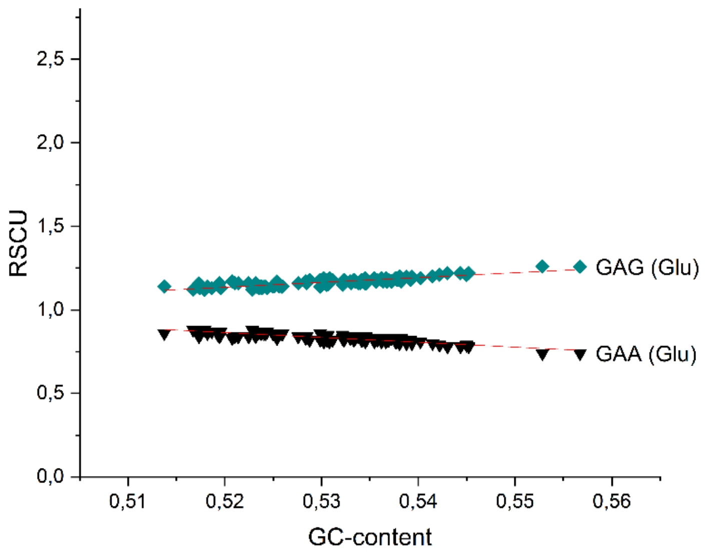


**Supplementary figure 7.a.** Impact of the GC content of coding genomes of mammalian species on their relative synonymous codon usage (RSCU) of the two **glutamic acid** codons.

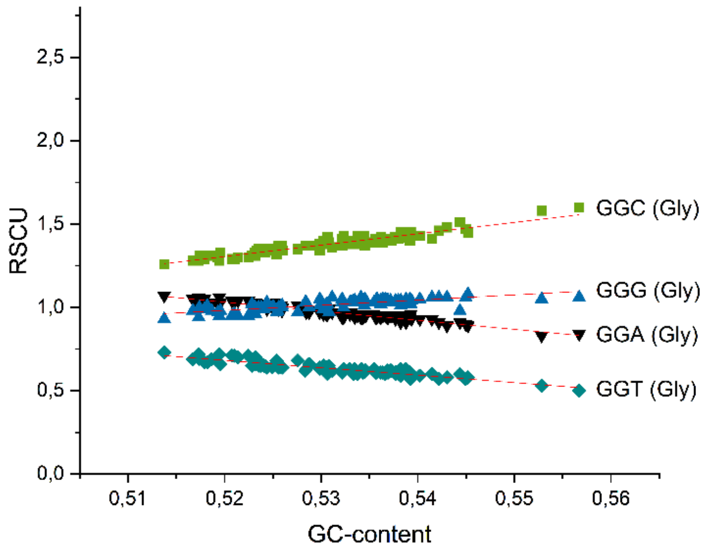


**Supplementary figure 8.a.** Impact of the GC content of coding genomes of mammalian species on their relative synonymous codon usage (RSCU) of the four **glycine** codons.

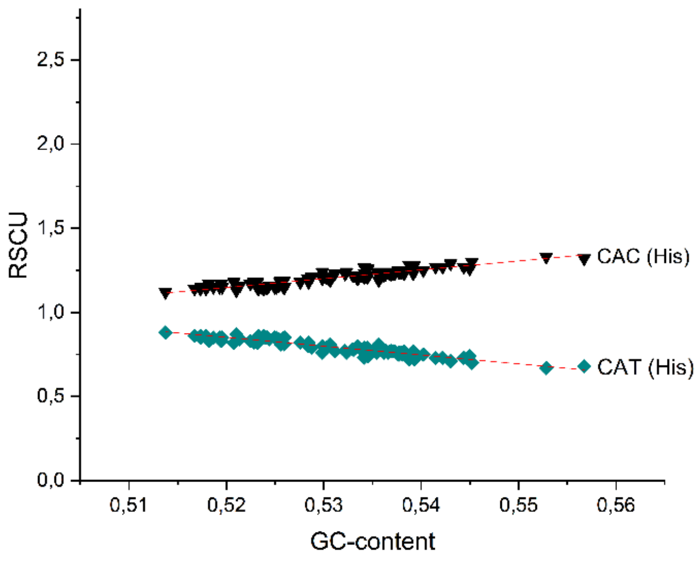


**Supplementary figure 9.a.** Impact of the GC content of coding genomes of mammalian species on their relative synonymous codon usage (RSCU) of the two **histidine** codons.

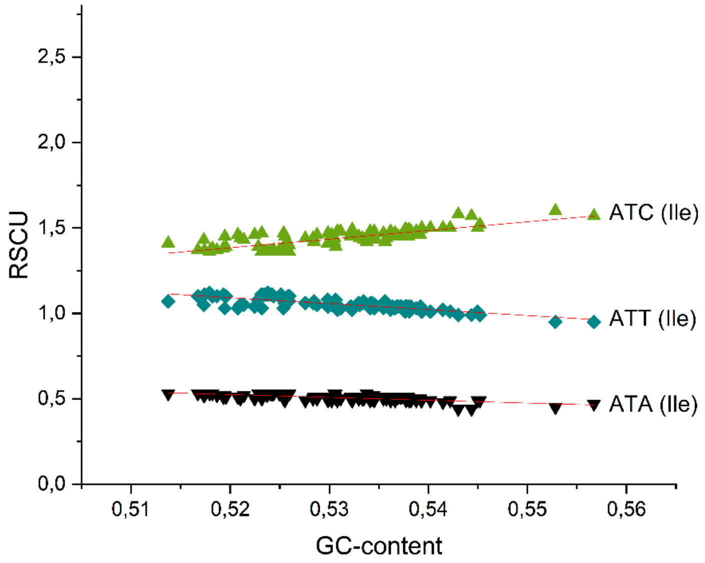


**Supplementary figure 10.a.** Impact of the GC content of coding genomes of mammalian species on their relative synonymous codon usage (RSCU) of the three **isoleucine** codons.

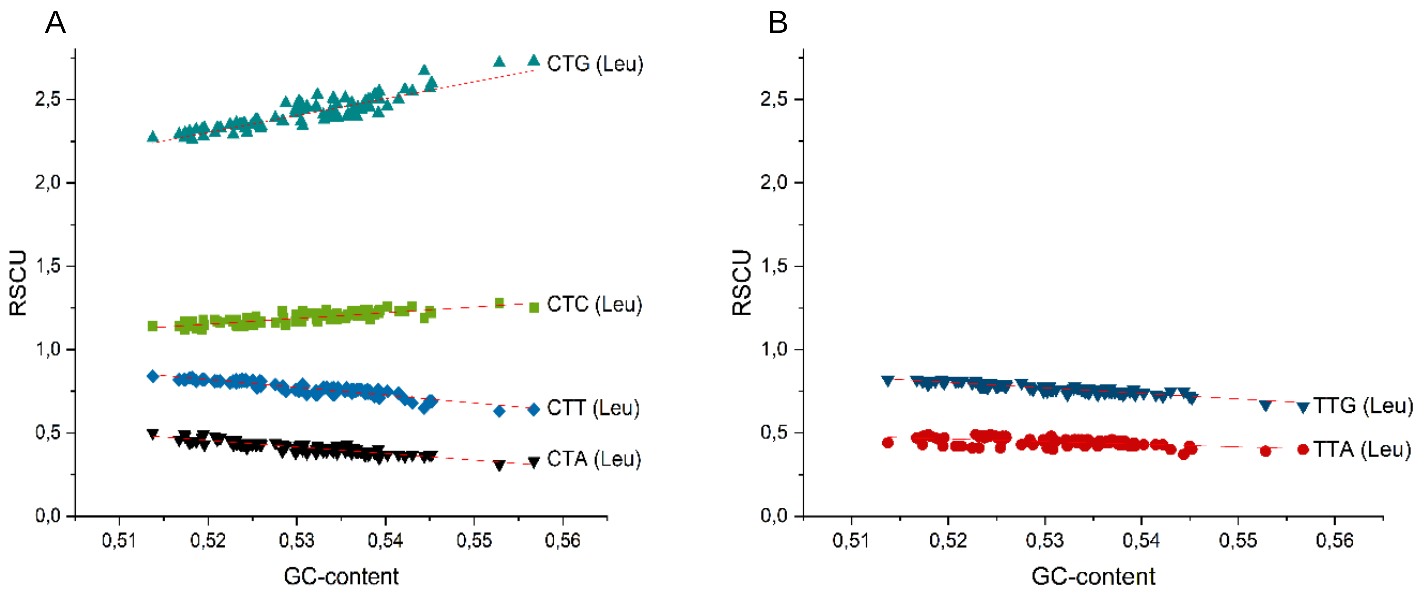


**Supplementary figure 11.a.** Impact of the GC content of coding genomes of mammalian species on their relative synonymous codon usage (RSCU) of the six **leucine** codons.

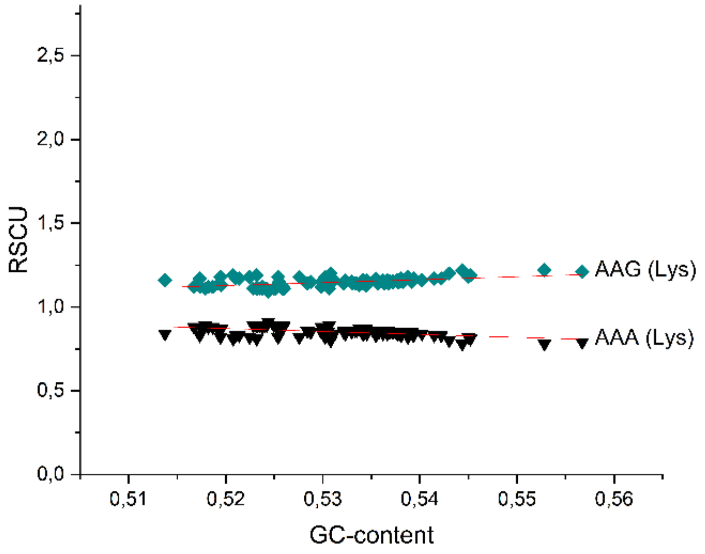


**Supplementary figure 12.a.** Impact of the GC content of coding genomes of mammalian species on their relative synonymous codon usage (RSCU) of the two **lysine** codons.

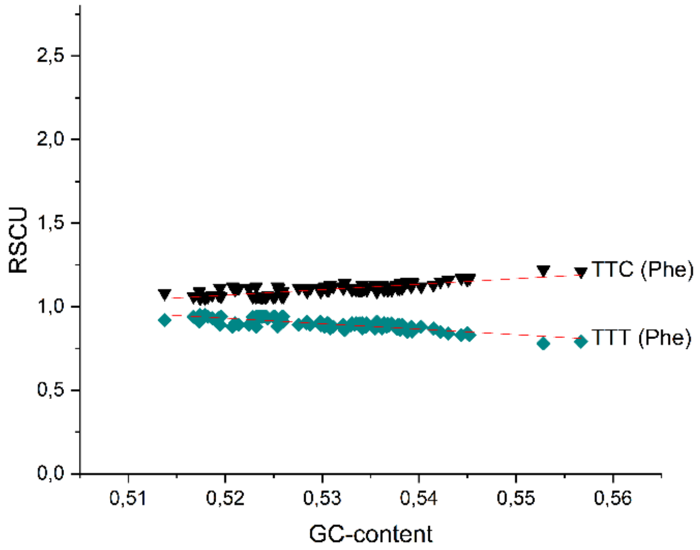


**Supplementary figure 13.a.** Impact of the GC content of coding genomes of mammalian species on their relative synonymous codon usage (RSCU) of the two **phenylalanine** codons.

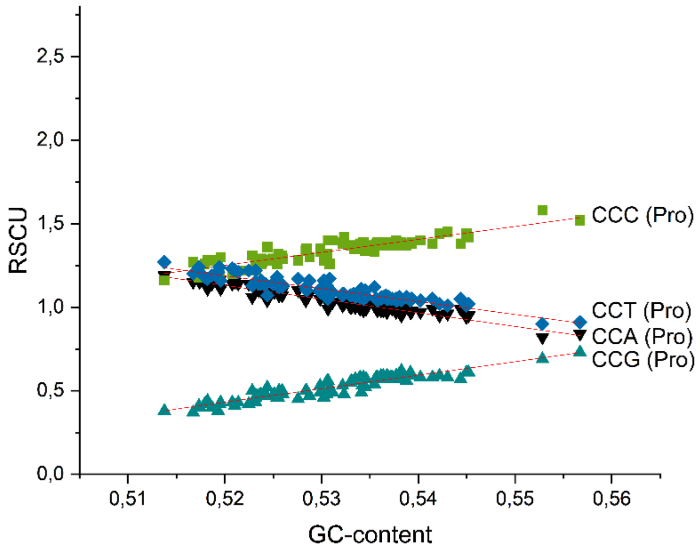


**Supplementary figure 14.a.** Impact of the GC content of coding genomes of mammalian species on their relative synonymous codon usage (RSCU) of the four **proline** codons.

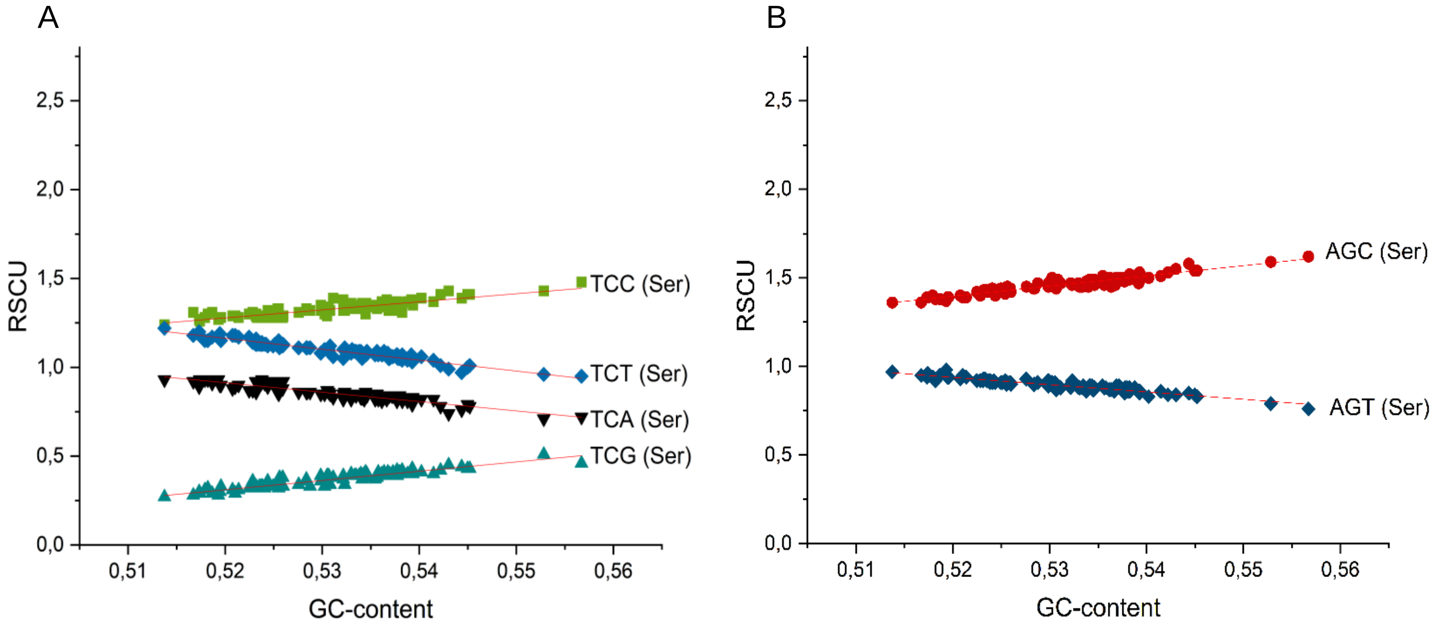


**Supplementary figure 15.a.** Impact of the GC content of coding genomes of mammalian species on their relative synonymous codon usage (RSCU) of the six **serine** codons.

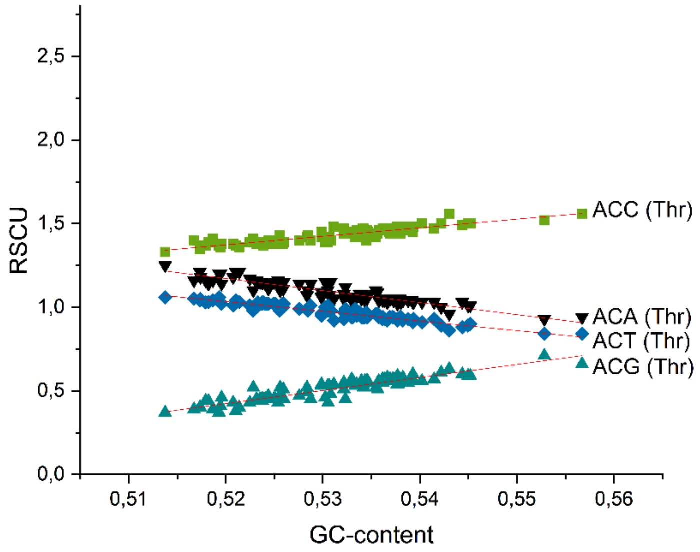


**Supplementary figure 16.a.** Impact of the GC content of coding genomes of mammalian species on their relative synonymous codon usage (RSCU) of the four **threonine** codons.

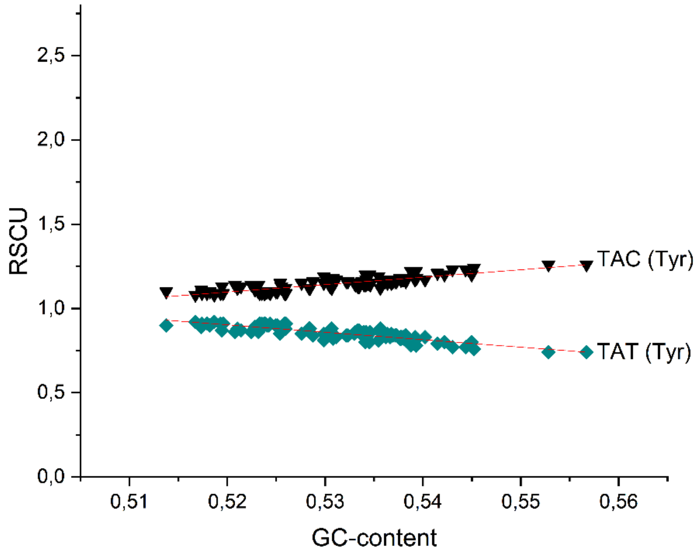


**Supplementary figure 17.a.** Impact of the GC content of coding genomes of mammalian species on their relative synonymous codon usage (RSCU) of the two **tyrosine** codons.

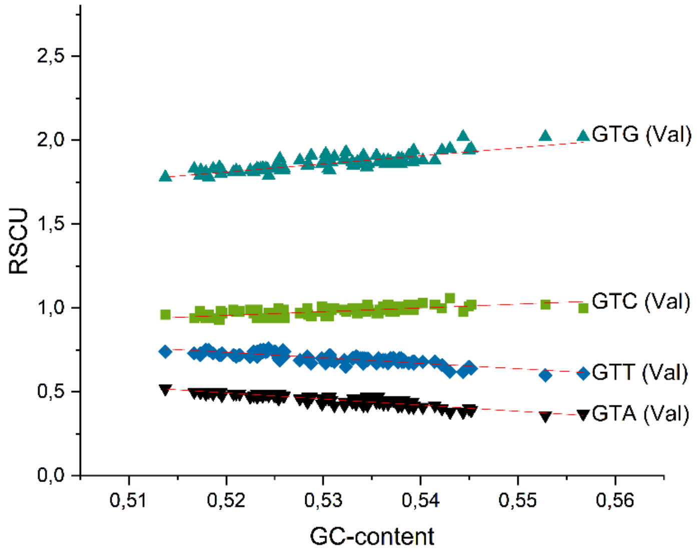


**Supplementary figure 18.a.** Impact of the GC content of coding genomes of mammalian species on their relative synonymous codon usage (RSCU) of the four **valine** codons.

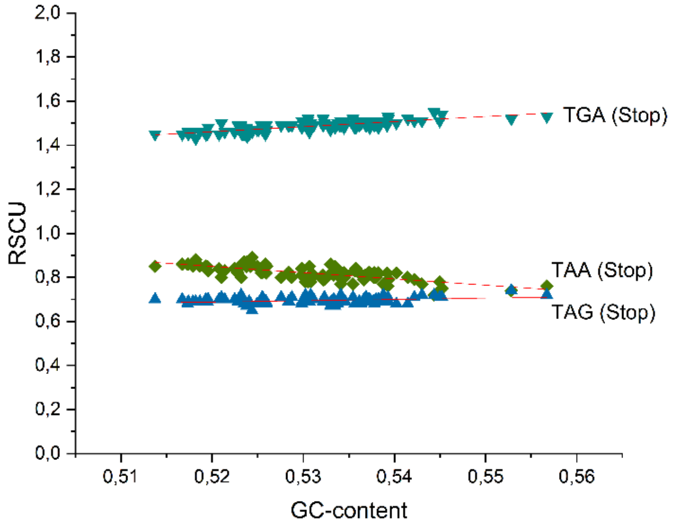


**Supplementary figure 19.a.** Impact of the GC content of coding genomes of mammalian species on their relative synonymous codon usage (RSCU) of the three **stop** codons.
