## Supplementary file 3 for "Codon bias coevolves with longevity"

### Supplementary file 3
#### Changes in codon usage and evolution of longevity in mammalian species

The file presents the plots reflecting changes in codon usage as a function of the lifespan of mammalian species. Note that in these analyses we have subtracted the RSCU values expected at the given GC content ($RSCU_{exp}$) from the actual RSCU values ($RSCU_{obs}$) and plotted these deviations as a function of the known lifespan (years) of the species. The parameters of the regression analyses of these plots are presented in Supplementary file 2.

The plots are presented in alphabetical order of the names of the amino acids; data for amino acids Met and Trp that are encoded by a single codon are not shown. We also present the data for the three stop codons at the end of the list of the 18 amino acids encoded by more than one synonymous codon.

In the case of the 18 amino acids and the termination codons the first figures (**Supplementary figures x.a**) present the plots reflecting changes in codon usage as a function of the lifespan of 19 rodent species. The second figures (**Supplementary figures x.b**) present the plots reflecting changes in codon usage as a function of the lifespan of the 96 mammalian species included in the present analysis.

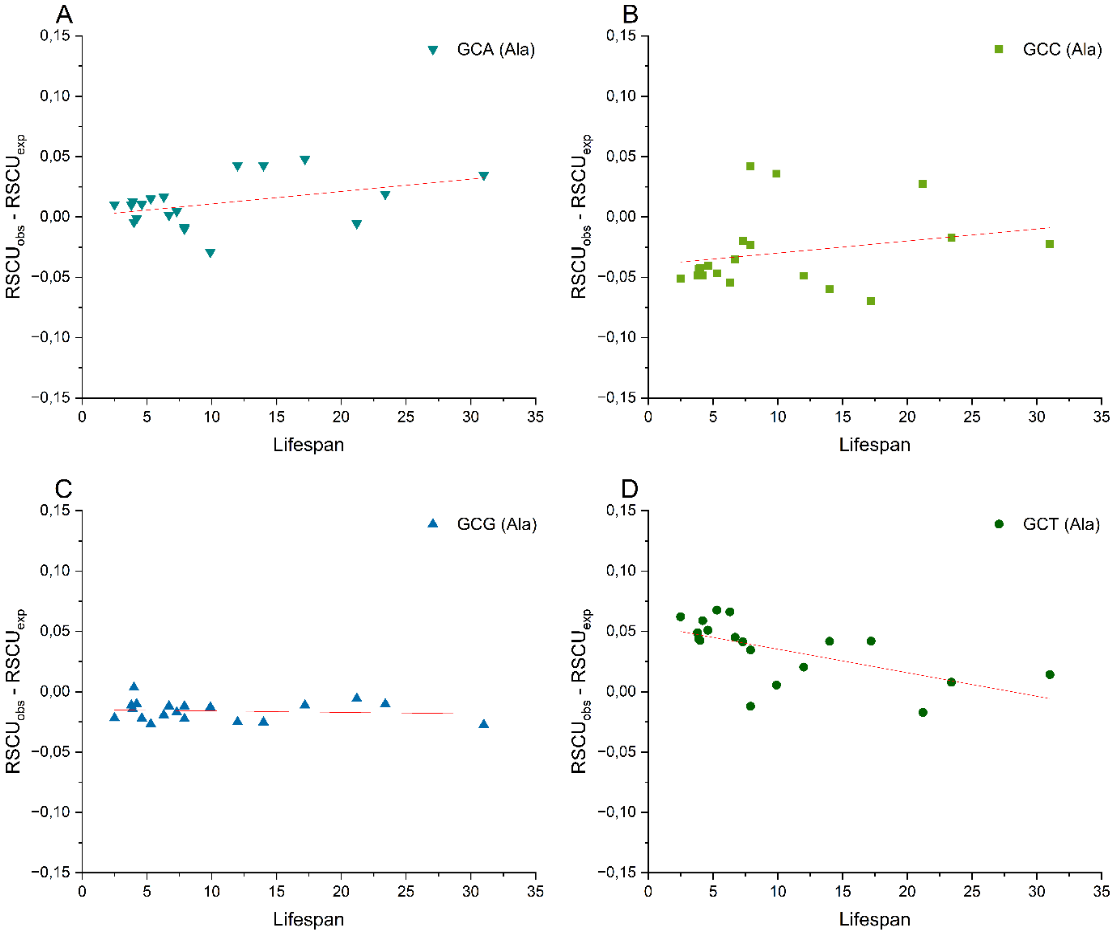


**Supplementary figure 1.a.** Differences between observed and expected relative synonymous codon usage (RSCU) values for codons encoding **alanine**, plotted against maximum lifespan (years) of 19 **rodent** species.

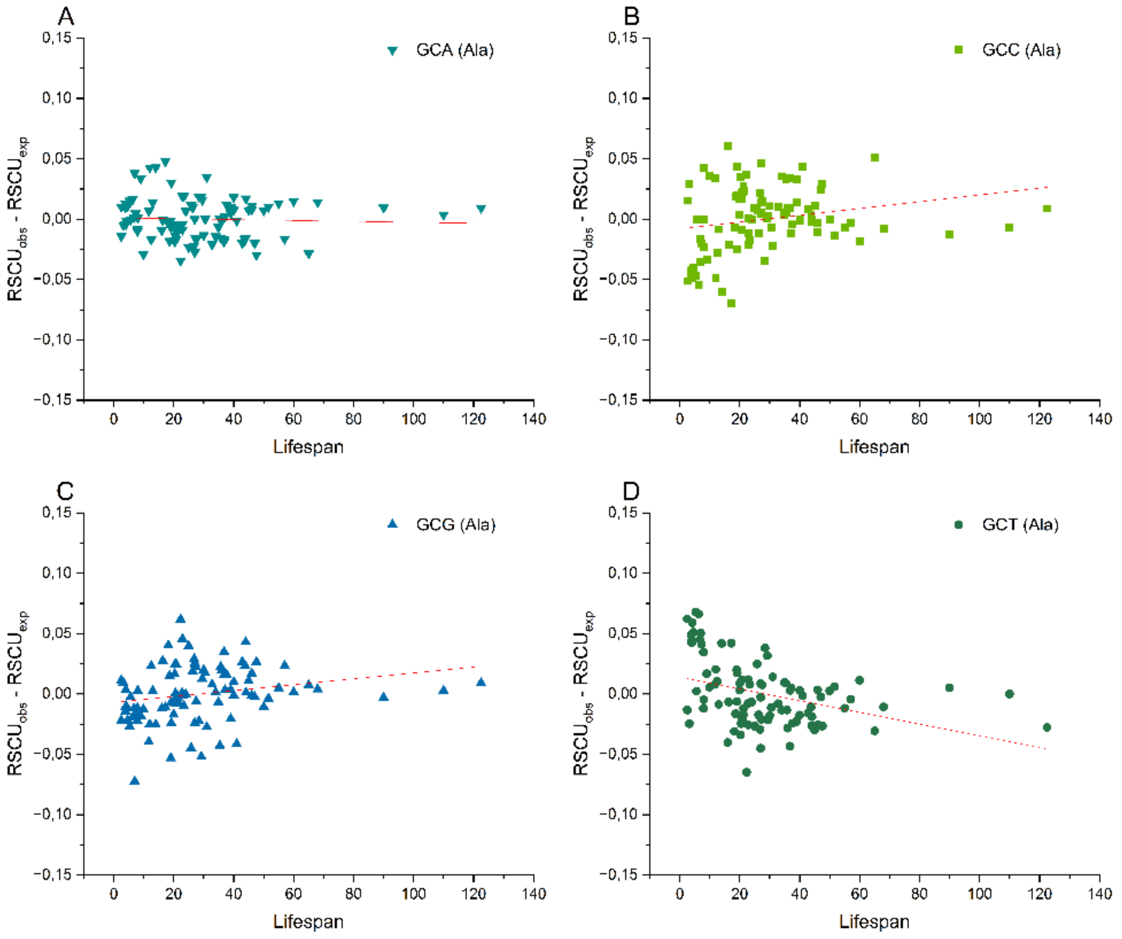


**Supplementary figure 1.b.** Differences between observed and expected relative synonymous codon usage (RSCU) values for codons encoding **alanine**, plotted against maximum lifespan (years) of 96 **mammalian** species.

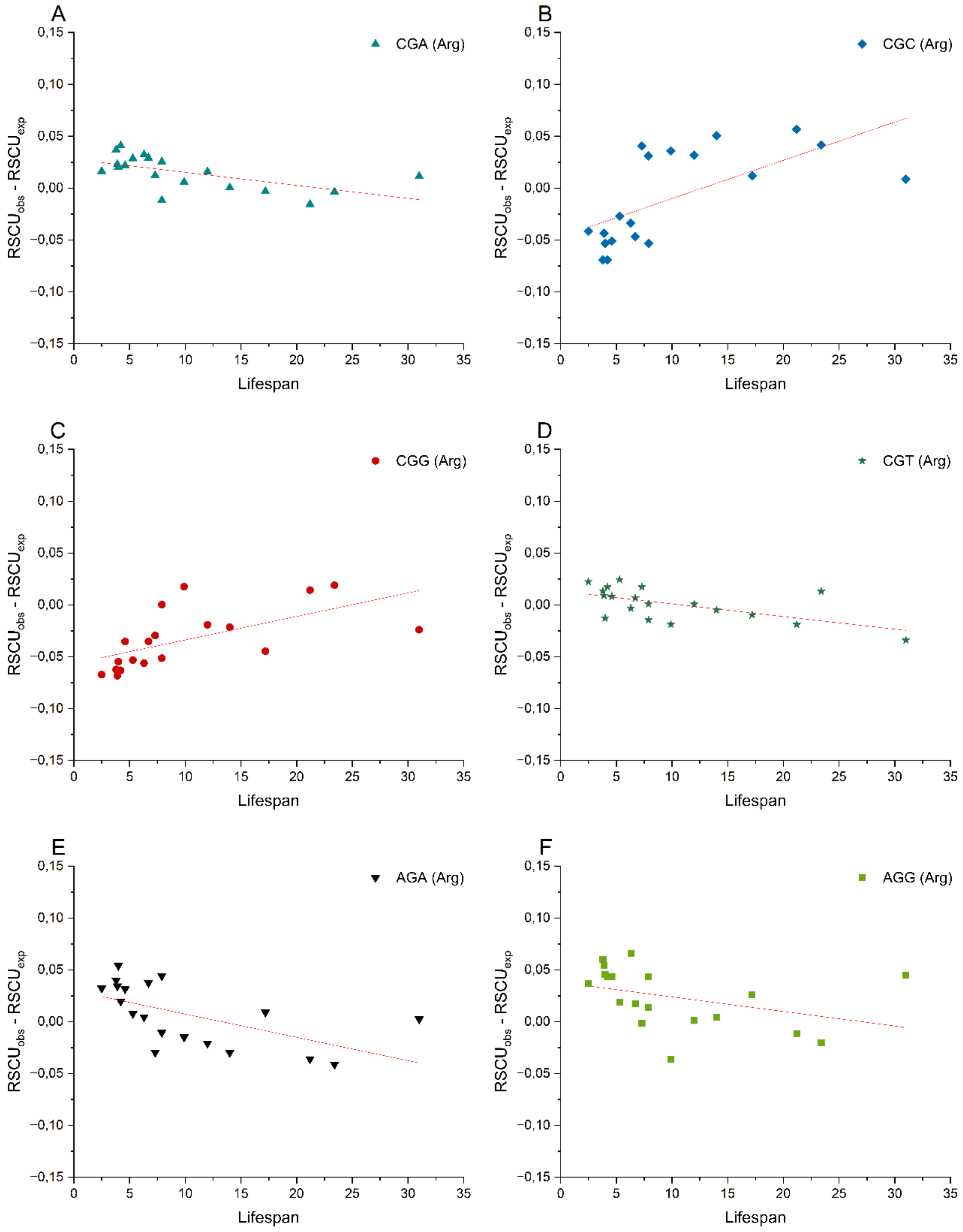


**Supplementary figure 2.a.** Differences between observed and expected relative synonymous codon usage (RSCU) values for codons encoding **arginine**, plotted against maximum lifespan (years) of 19 **rodent** species.

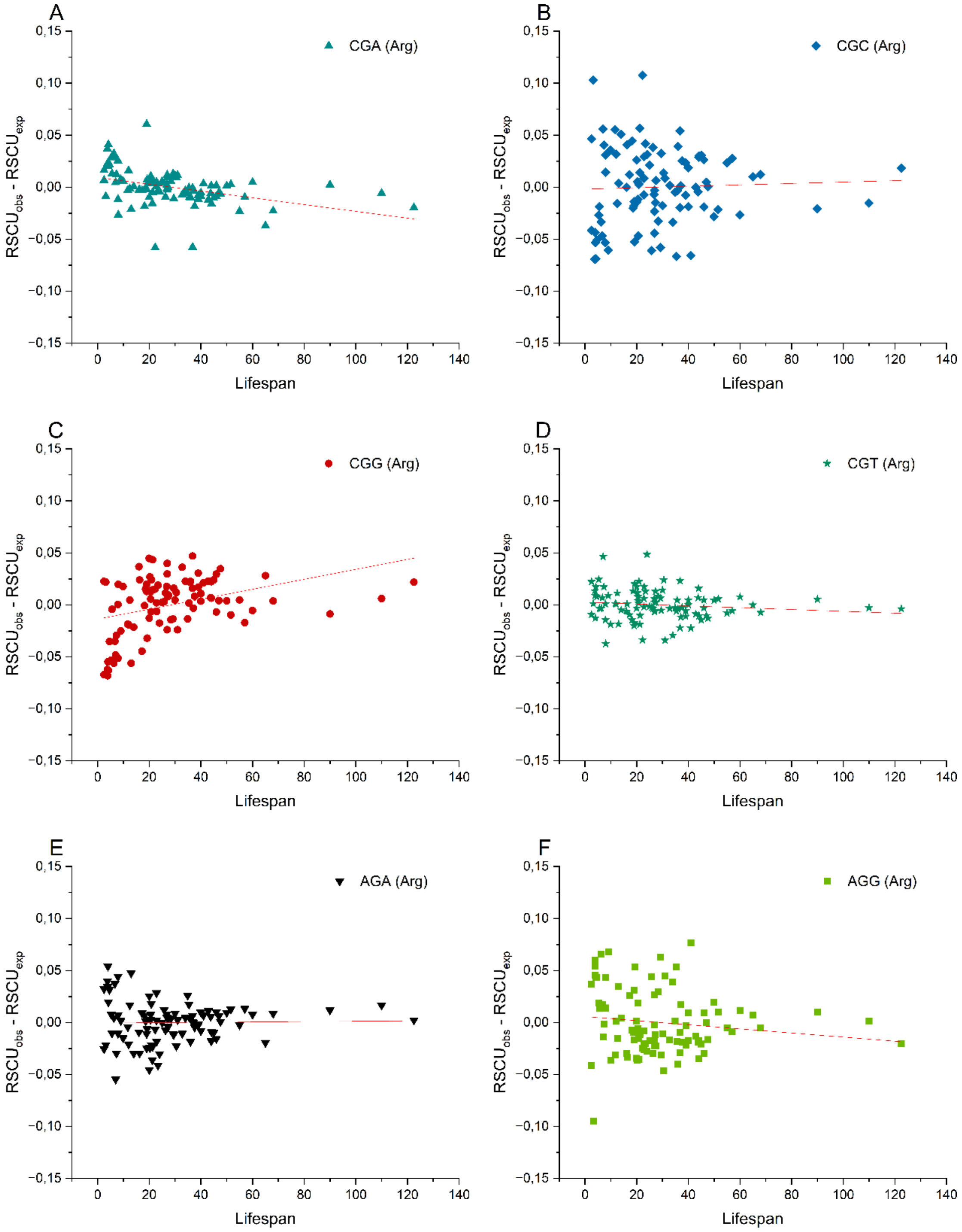


**Supplementary figure 2.b.** Differences between observed and expected relative synonymous codon usage (RSCU) values for codons encoding **arginine**, plotted against maximum lifespan (years) of 96 **mammalian** species.

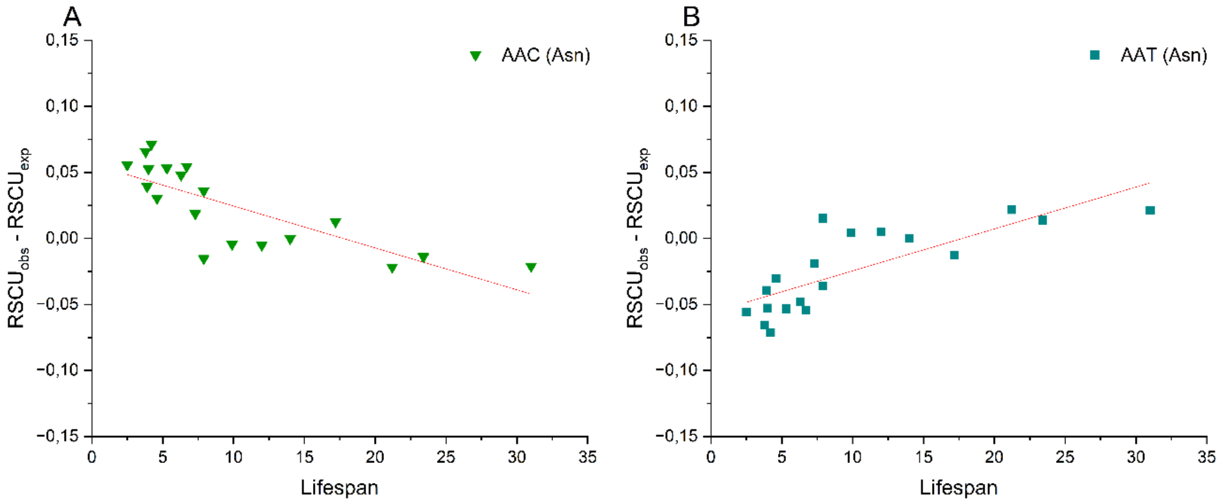


**Supplementary figure 3.a.** Differences between observed and expected relative synonymous codon usage (RSCU) values for codons encoding **asparagine**, plotted against maximum lifespan (years) of 19 **rodent** species.

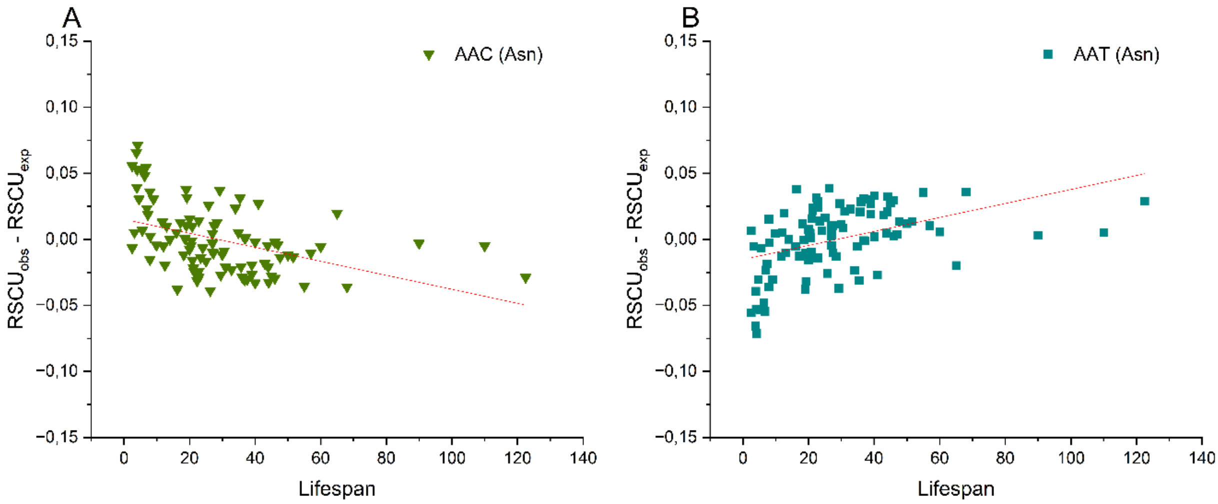


**Supplementary figure 3.b.** Differences between observed and expected relative synonymous codon usage (RSCU) values for codons encoding **asparagine**, plotted against maximum lifespan (years) of 96 **mammalian** species.

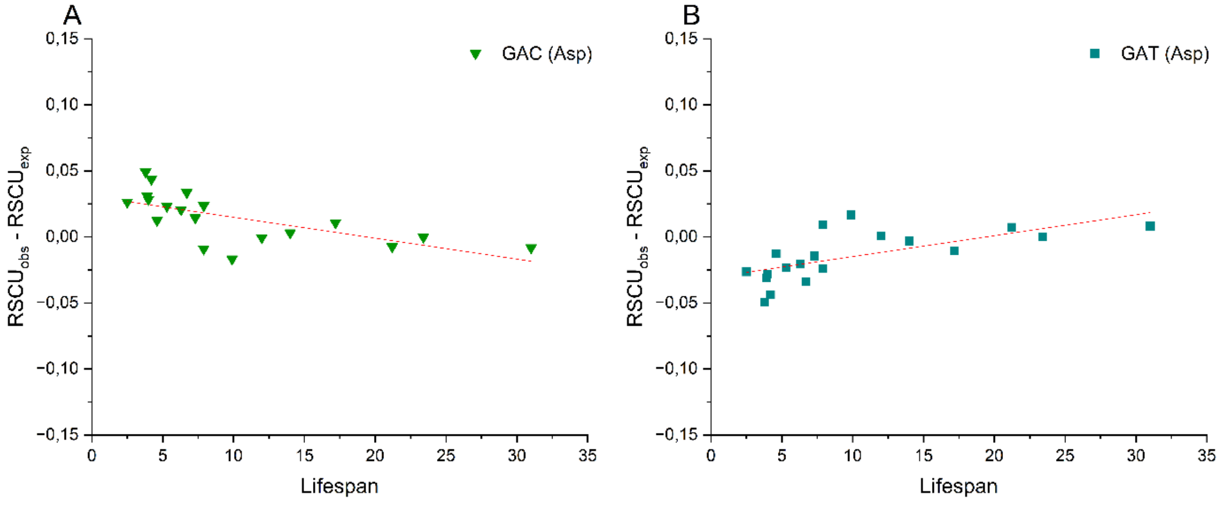


**Supplementary figure 4.a.** Differences between observed and expected relative synonymous codon usage (RSCU) values for codons encoding **aspartic acid**, plotted against maximum lifespan (years) of 19 **rodent** species.

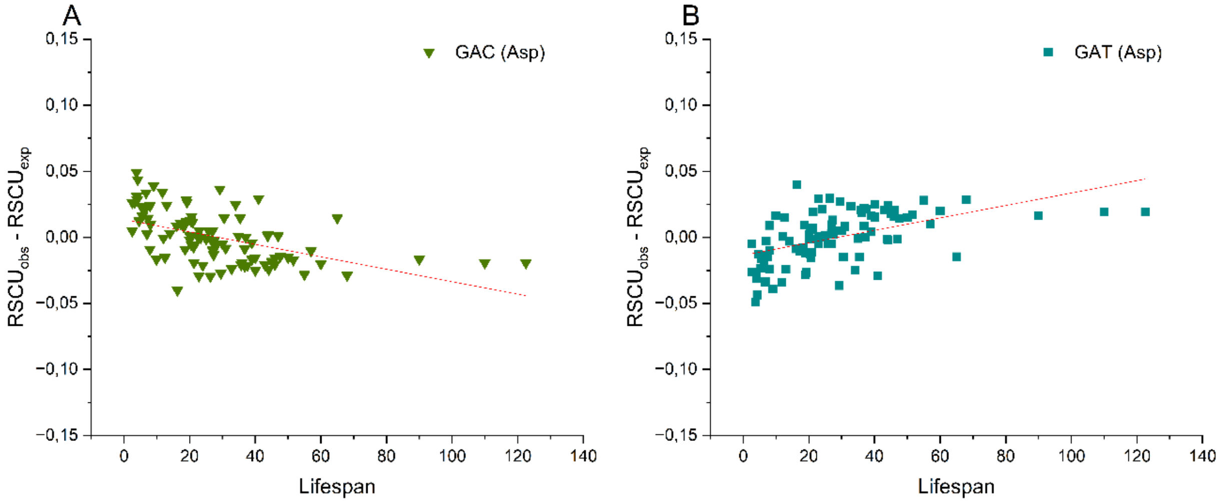


**Supplementary figure 4.b.** Differences between observed and expected relative synonymous codon usage (RSCU) values for codons encoding **aspartic acid**, plotted against maximum lifespan (years) of 96 **mammalian** species.

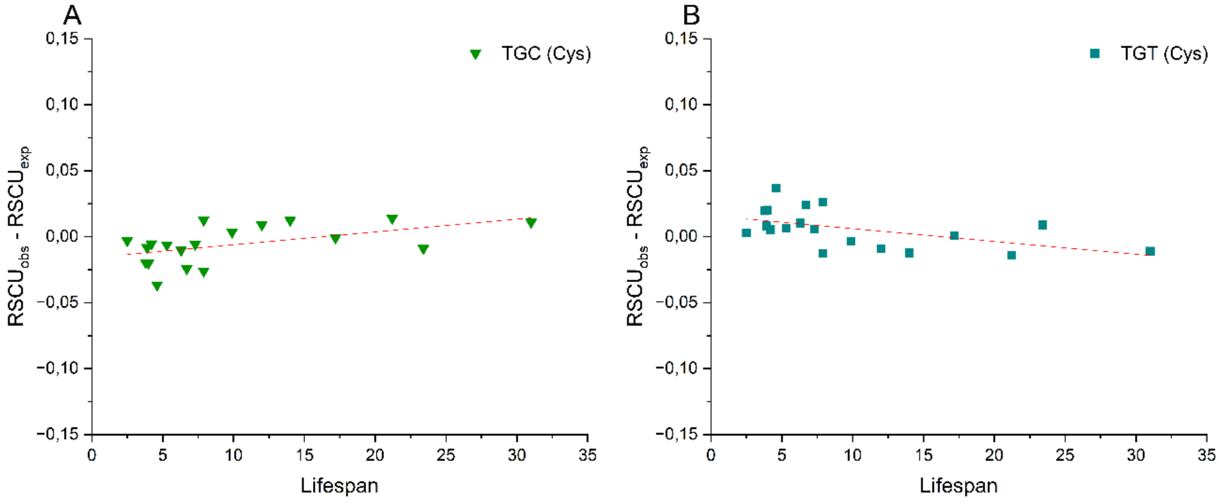


**Supplementary figure 5.a.** Differences between observed and expected relative synonymous codon usage (RSCU) values for codons encoding **cysteine**, plotted against maximum lifespan (years) of 19 **rodent** species.

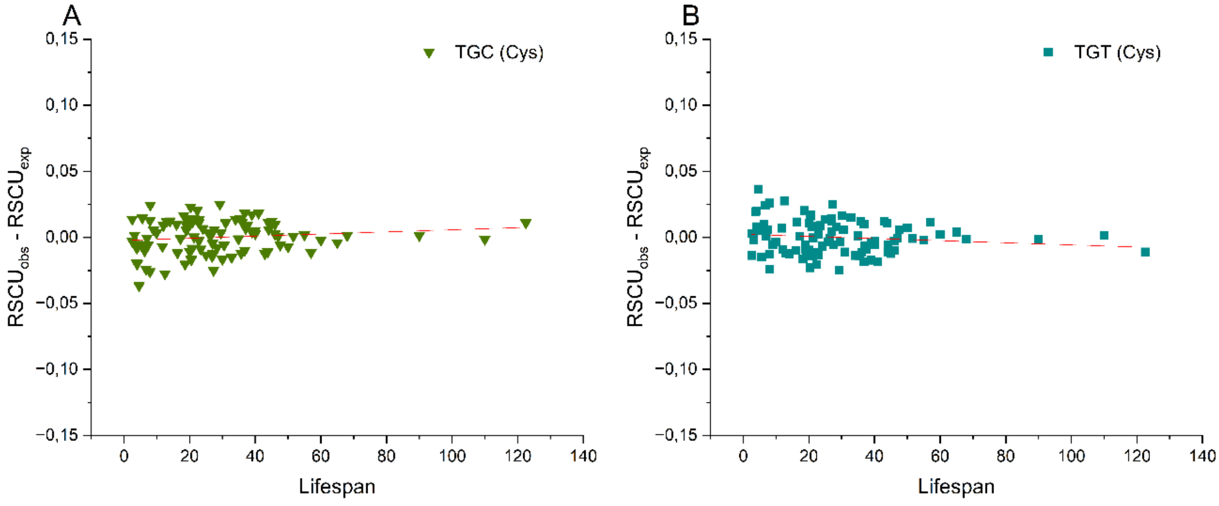


**Supplementary figure 5.b.** Differences between observed and expected relative synonymous codon usage (RSCU) values for codons encoding **cysteine**, plotted against maximum lifespan (years) of 96 **mammalian** species.

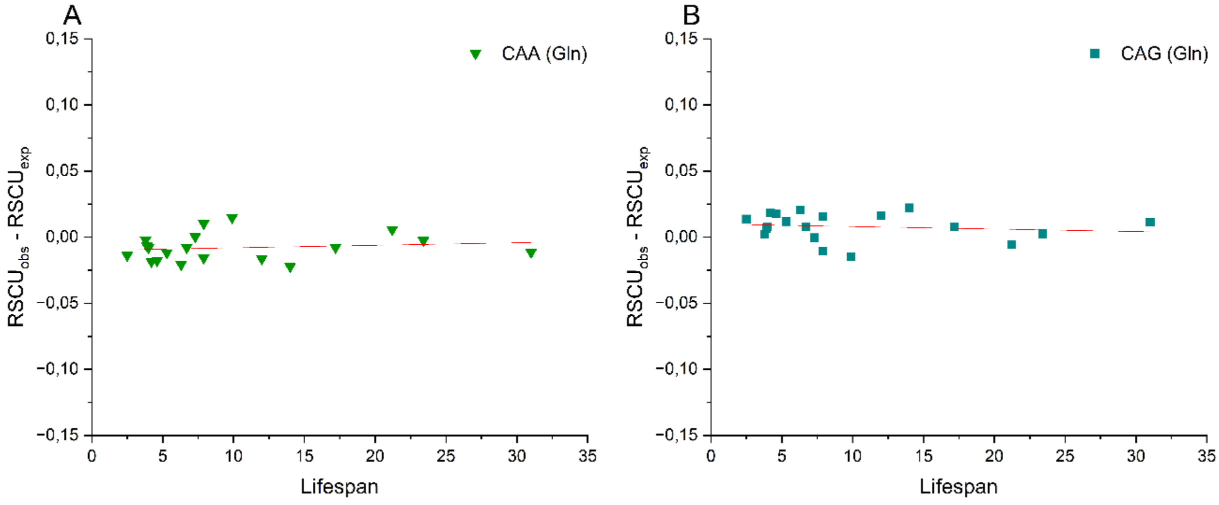


**Supplementary figure 6.a.** Differences between observed and expected relative synonymous codon usage (RSCU) values for codons encoding **glutamine**, plotted against maximum lifespan (years) of 19 **rodent** species.

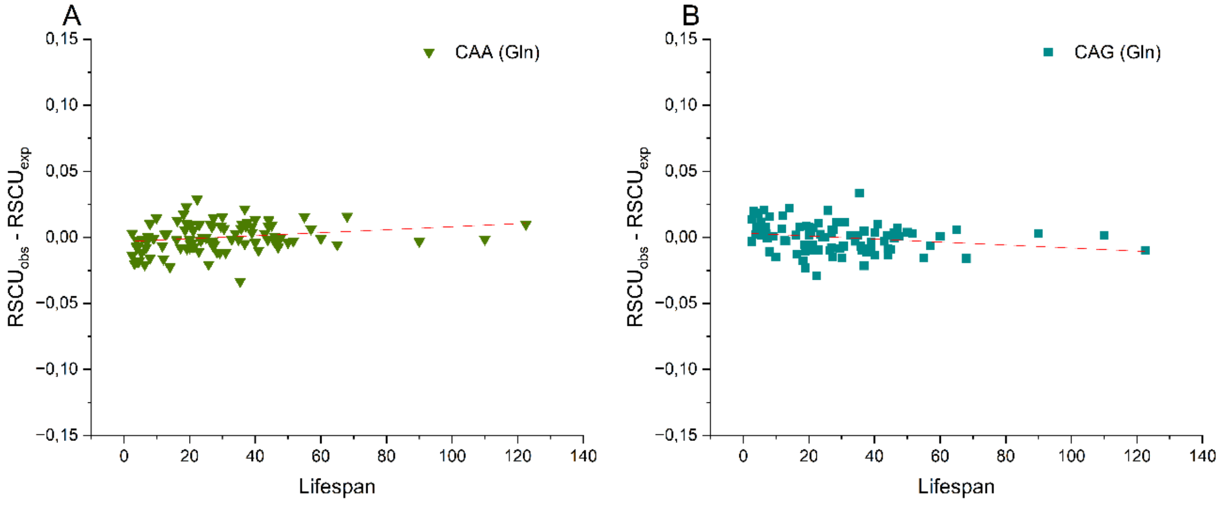


**Supplementary figure 6.b.** Differences between observed and expected relative synonymous codon usage (RSCU) values for codons encoding **glutamine**, plotted against maximum lifespan (years) of 96 **mammalian** species.

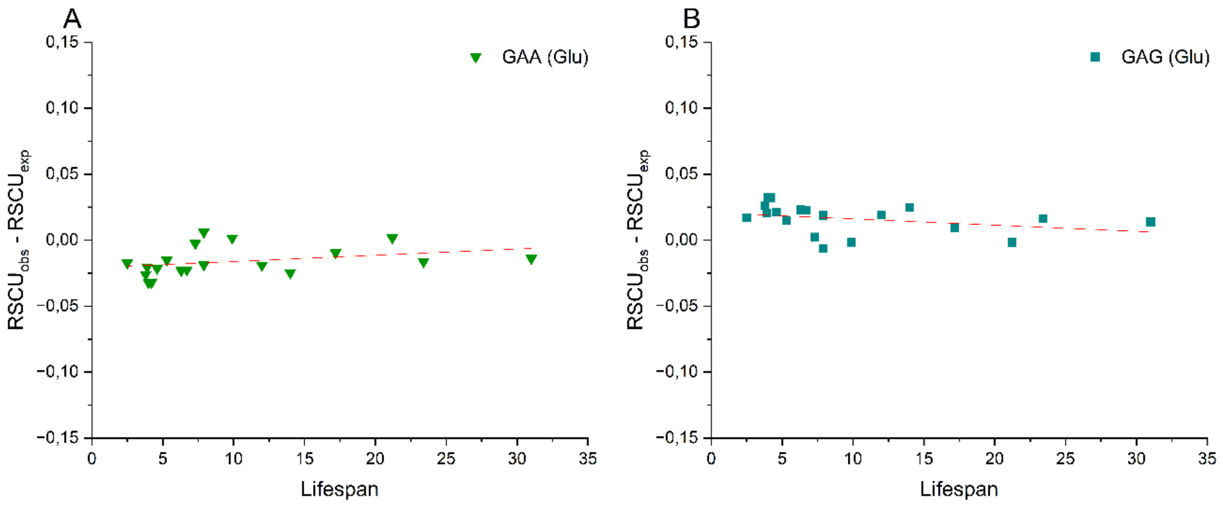


**Supplementary figure 7.a.** Differences between observed and expected relative synonymous codon usage (RSCU) values for codons encoding **glutamic acid**, plotted against maximum lifespan (years) of 19 **rodent** species.

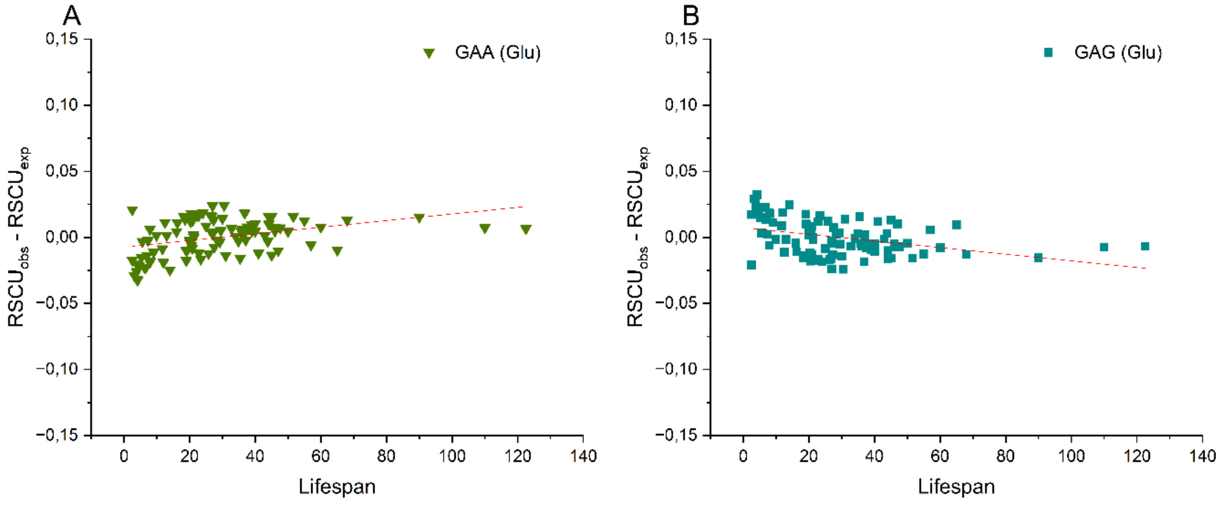


**Supplementary figure 7.b.** Differences between observed and expected relative synonymous codon usage (RSCU) values for codons encoding **glutamic acid**, plotted against maximum lifespan (years) of 96 **mammalian** species.

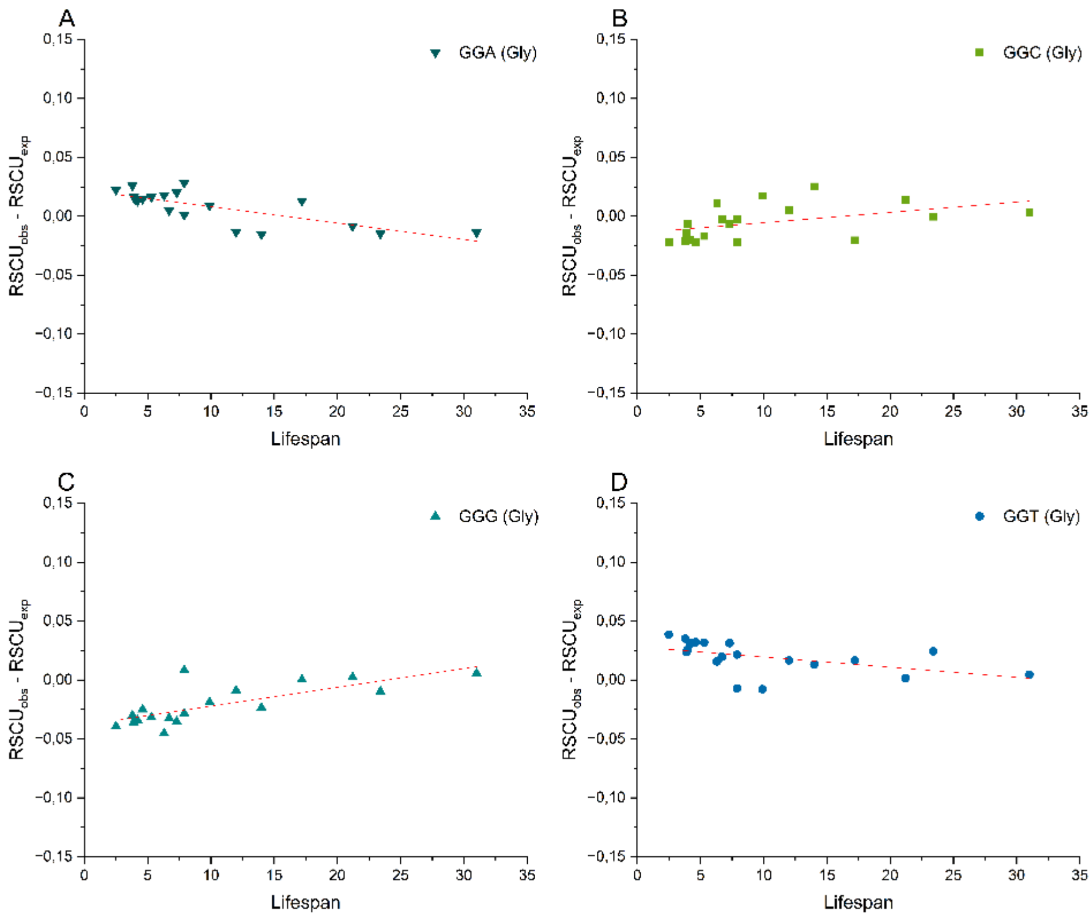


**Supplementary figure 8.a.** Differences between observed and expected relative synonymous codon usage (RSCU) values for codons encoding **glycine**, plotted against maximum lifespan (years) of 19 **rodent** species.

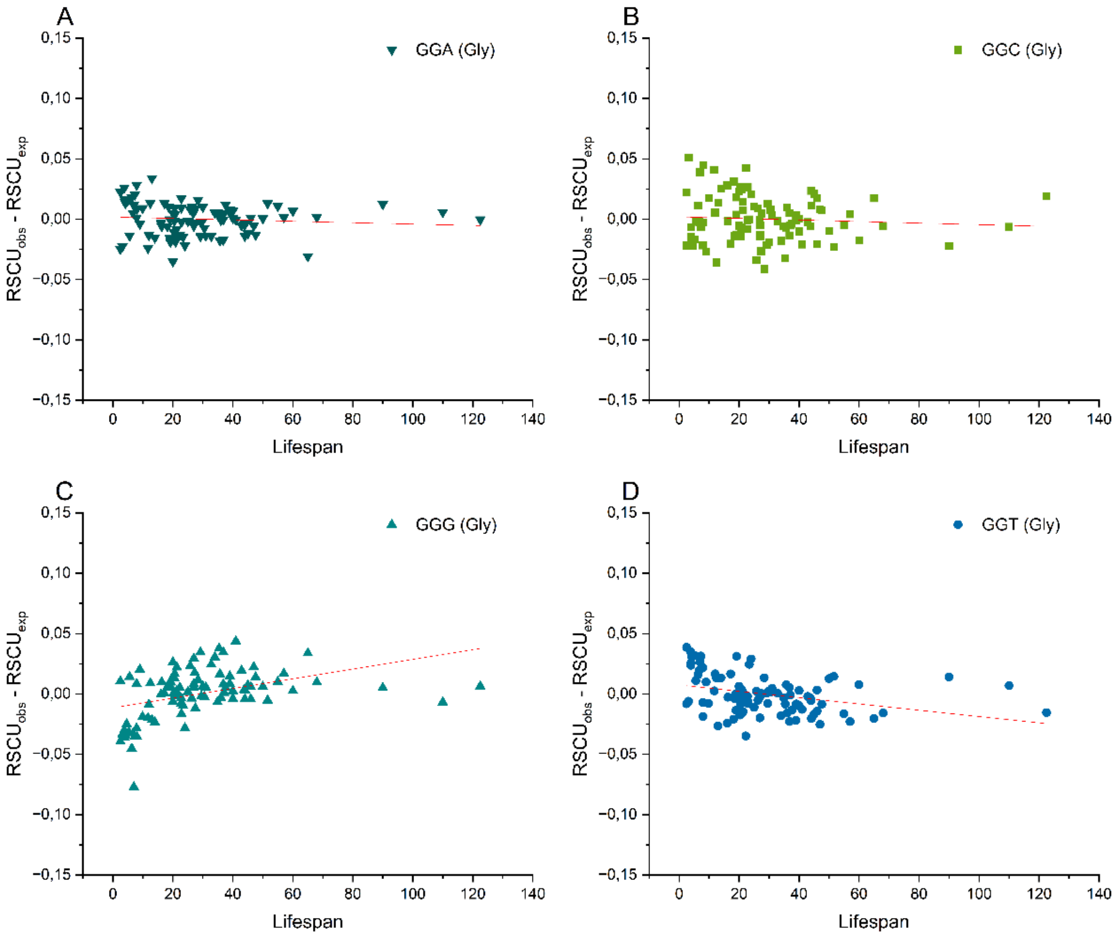


**Supplementary figure 8.b.** Differences between observed and expected relative synonymous codon usage (RSCU) values for codons encoding **glycine**, plotted against maximum lifespan (years) of 96 **mammalian** species.

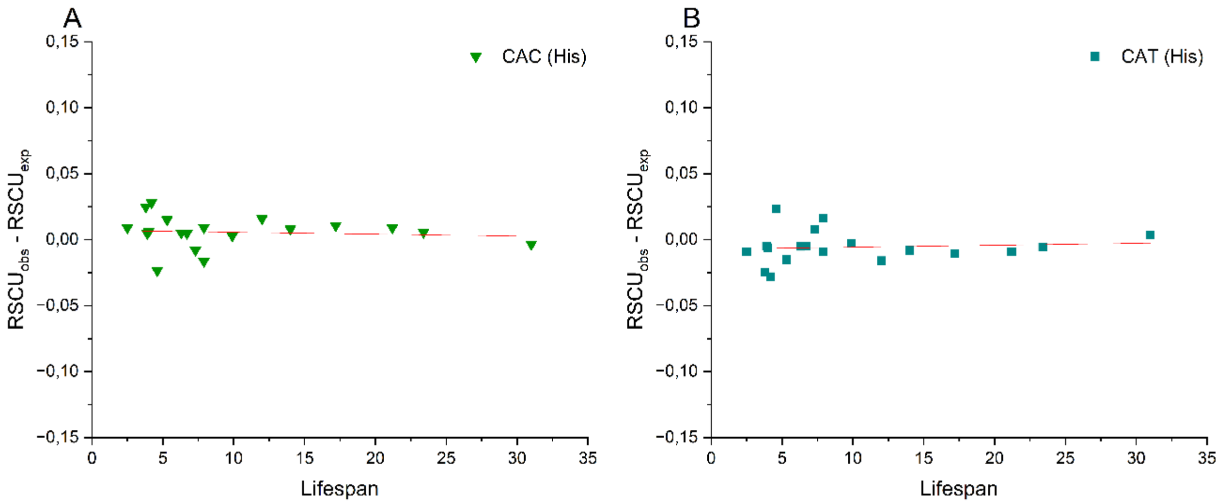


**Supplementary figure 9.a.** Differences between observed and expected relative synonymous codon usage (RSCU) values for codons encoding **histidine**, plotted against maximum lifespan (years) of 19 **rodent** species.

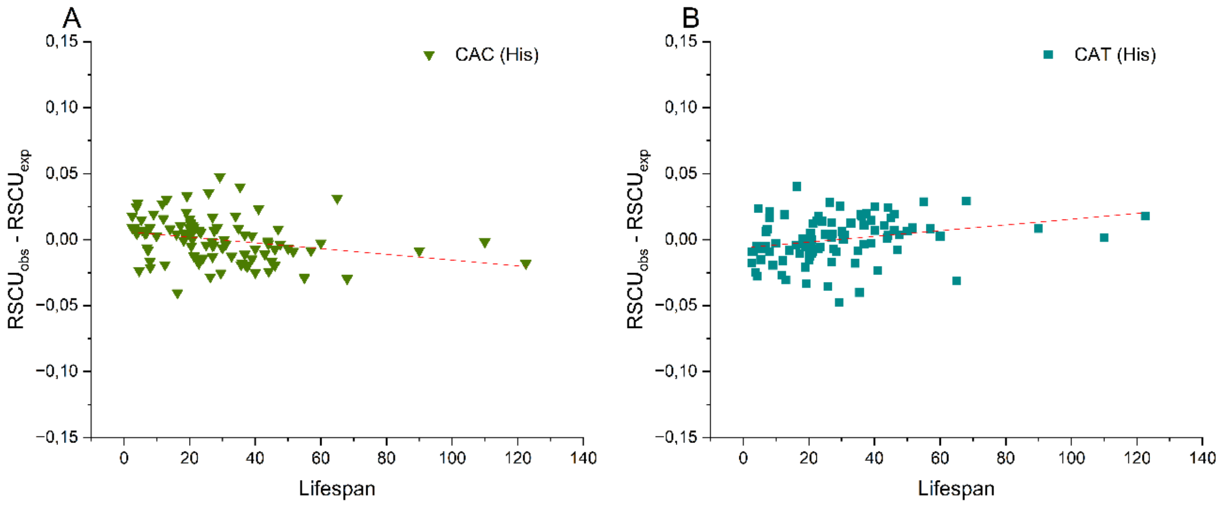


**Supplementary figure 9.b.** Differences between observed and expected relative synonymous codon usage (RSCU) values for codons encoding **histidine**, plotted against maximum lifespan (years) of 96 **mammalian** species.

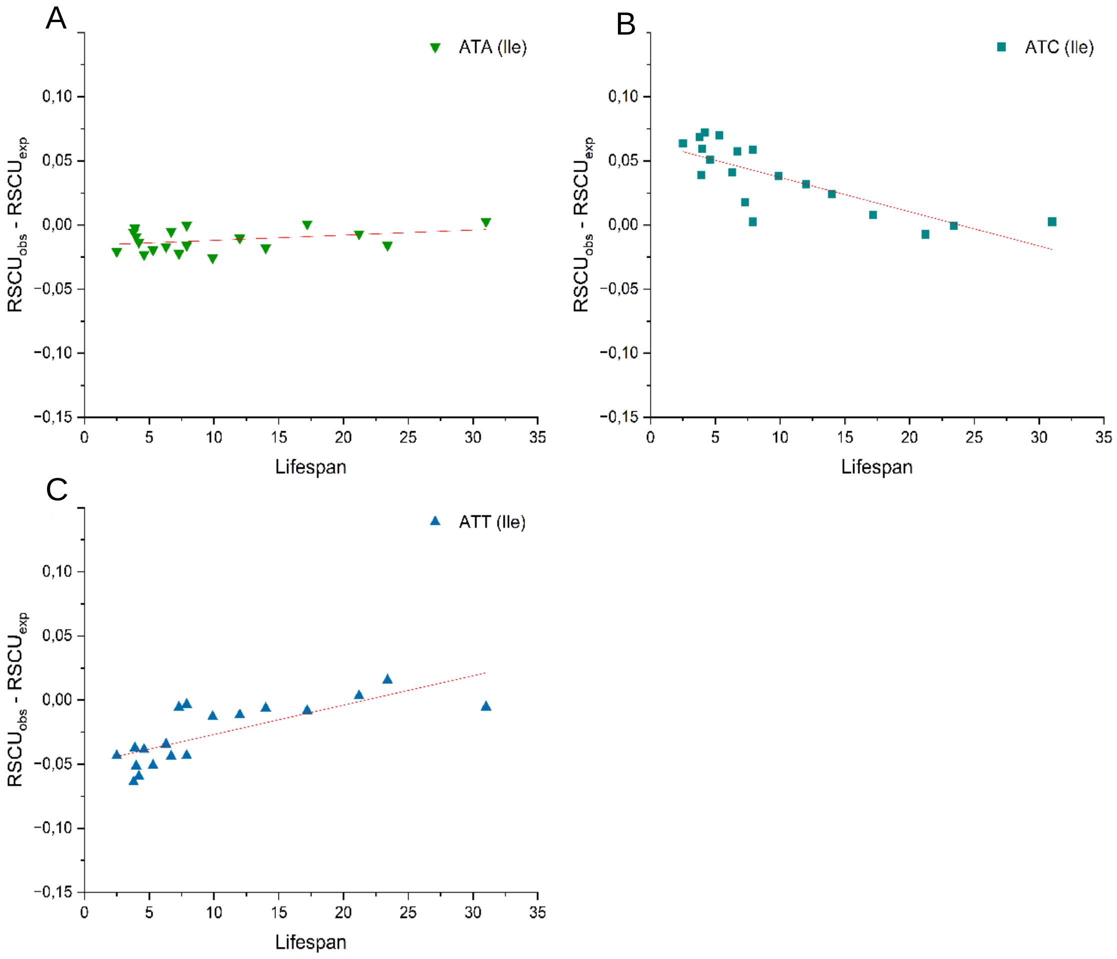


**Supplementary figure 10.a.** Differences between observed and expected relative synonymous codon usage (RSCU) values for codons encoding **isoleucine**, plotted against maximum lifespan (years) of 19 **rodent** species.

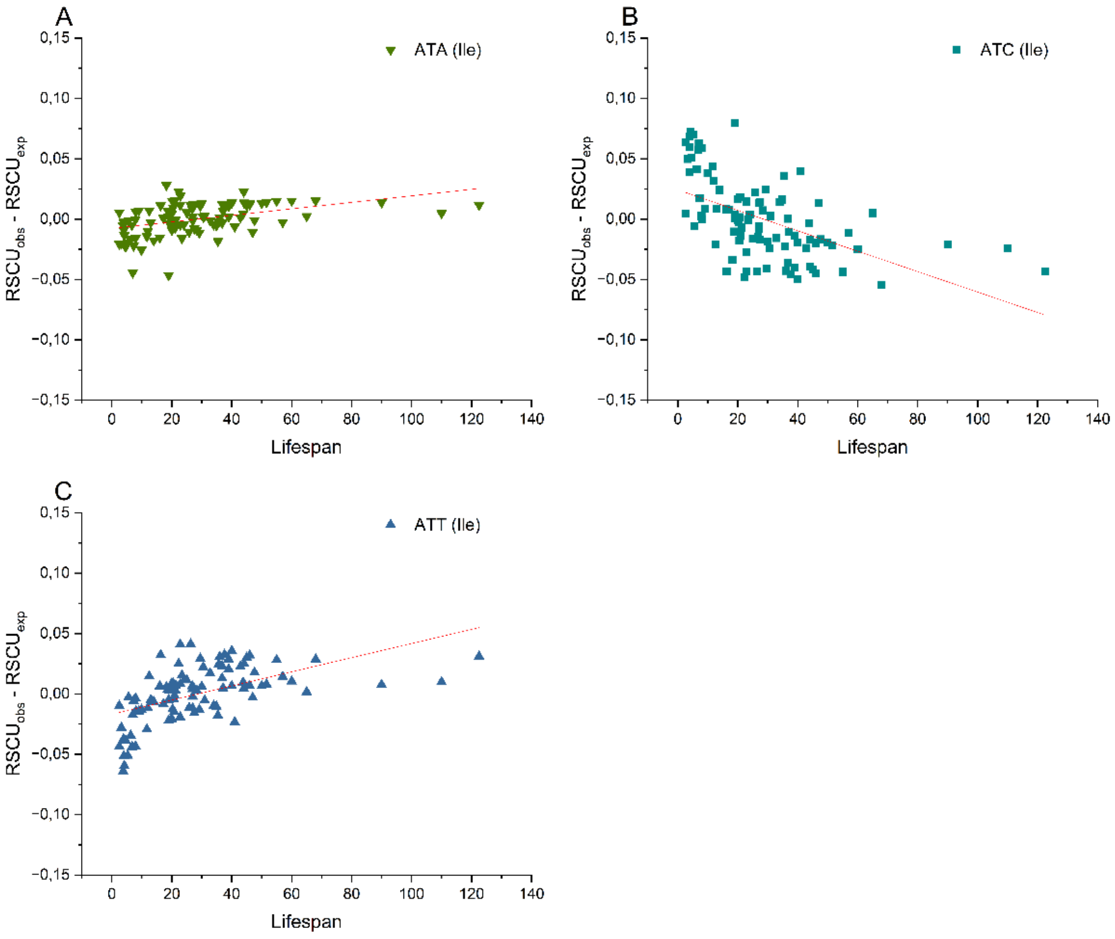


**Supplementary figure 10.b.** Differences between observed and expected relative synonymous codon usage (RSCU) values for codons encoding **isoleucine**, plotted against maximum lifespan (years) of 96 **mammalian** species.

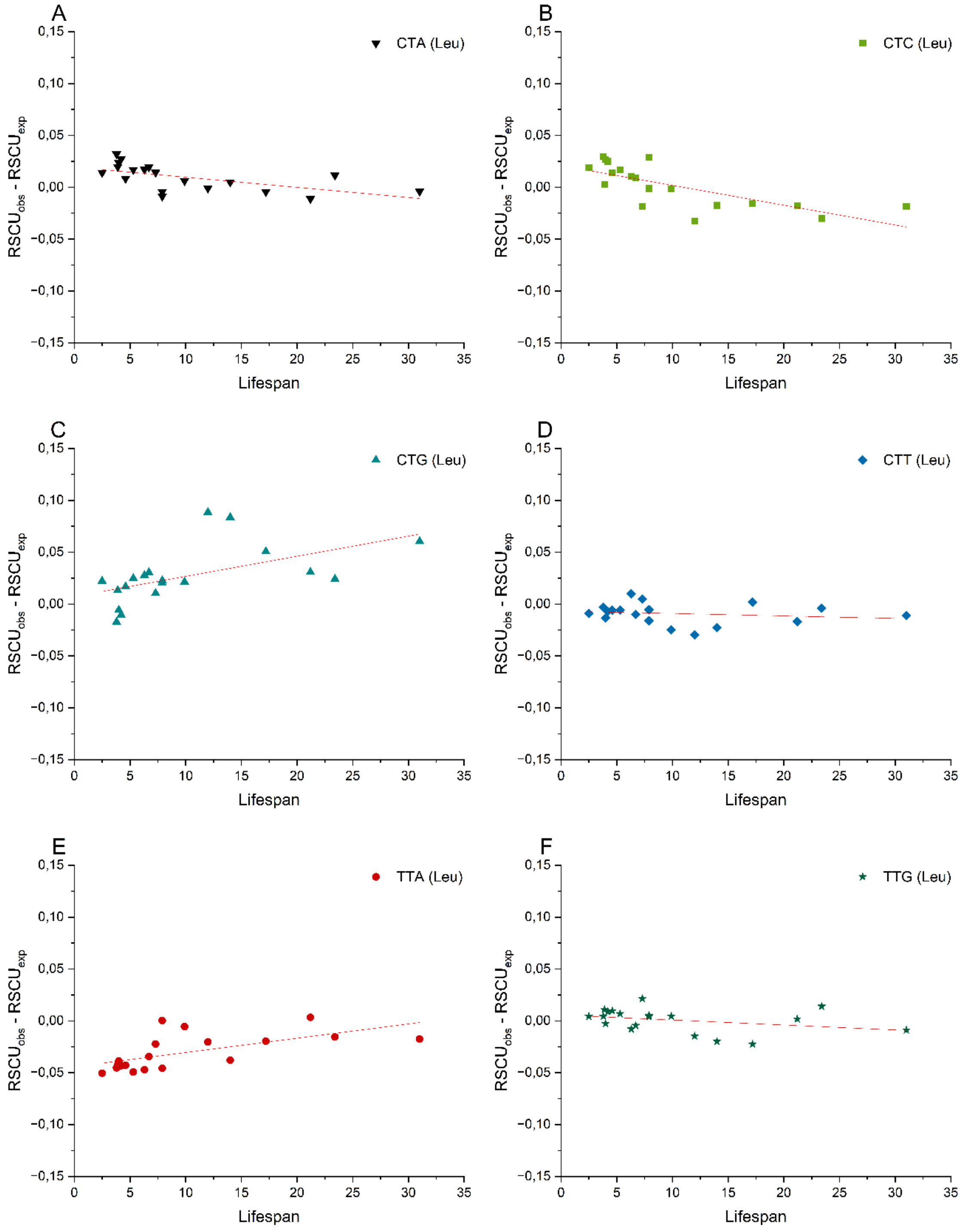


**Supplementary figure 11.a.** Differences between observed and expected relative synonymous codon usage (RSCU) values for codons encoding **leucine**, plotted against maximum lifespan (years) of 19 **rodent** species.

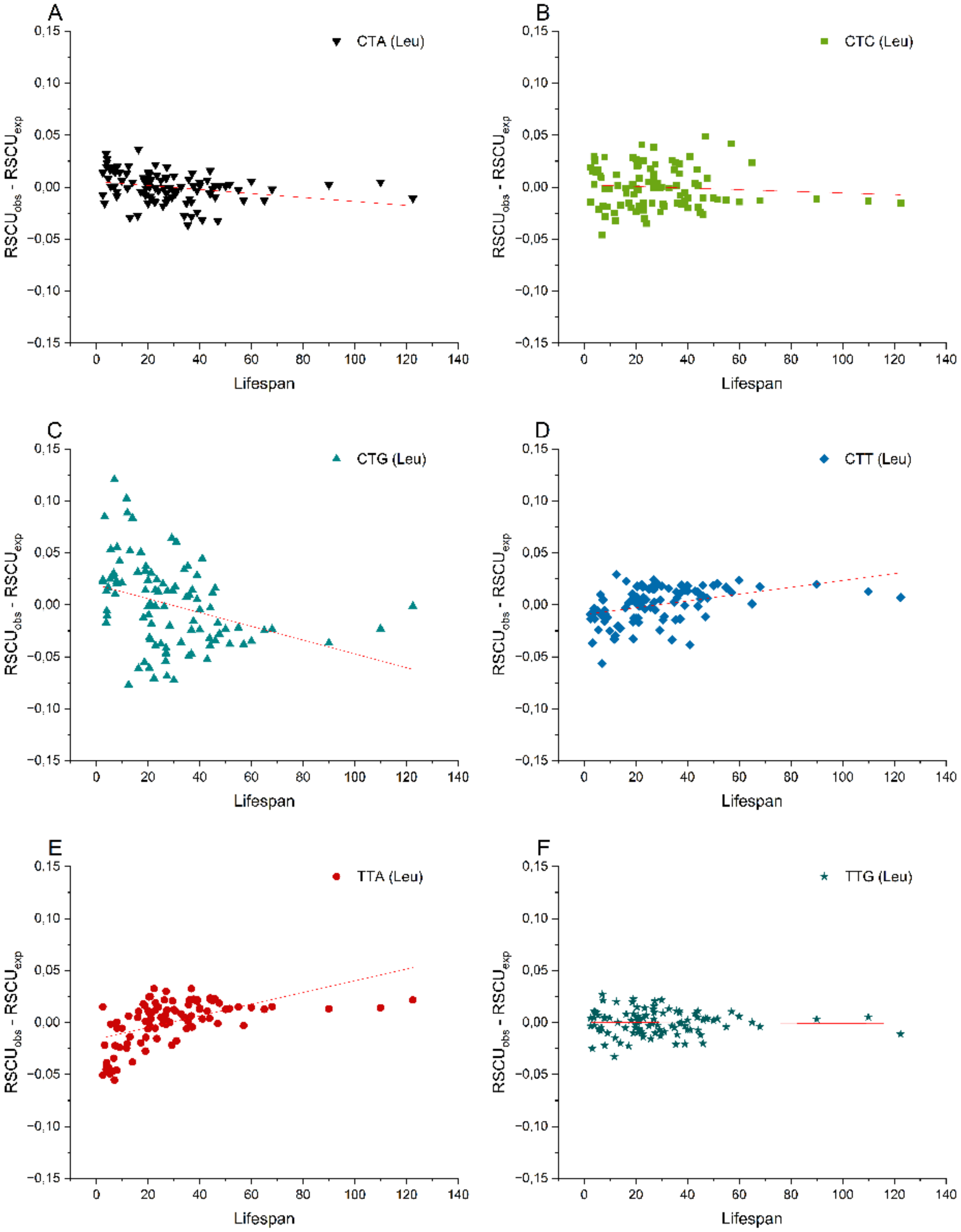


**Supplementary figure 11.b.** Differences between observed and expected relative synonymous codon usage (RSCU) values for codons encoding **leucine**, plotted against maximum lifespan (years) of 96 **mammalian** species.

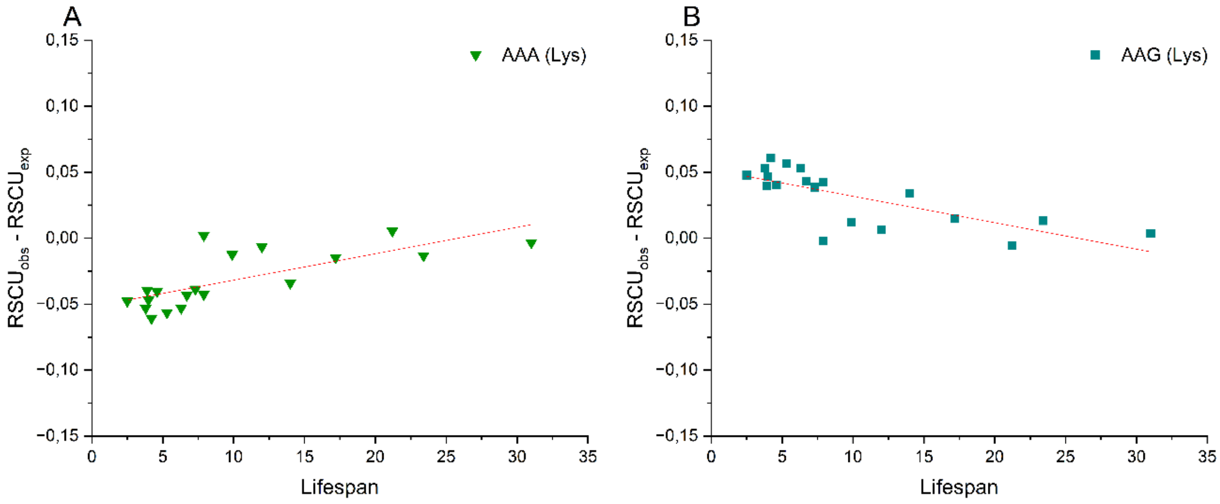


**Supplementary figure 12.a.** Differences between observed and expected relative synonymous codon usage (RSCU) values for codons encoding **lysine**, plotted against maximum lifespan (years) of 19 **rodent** species.

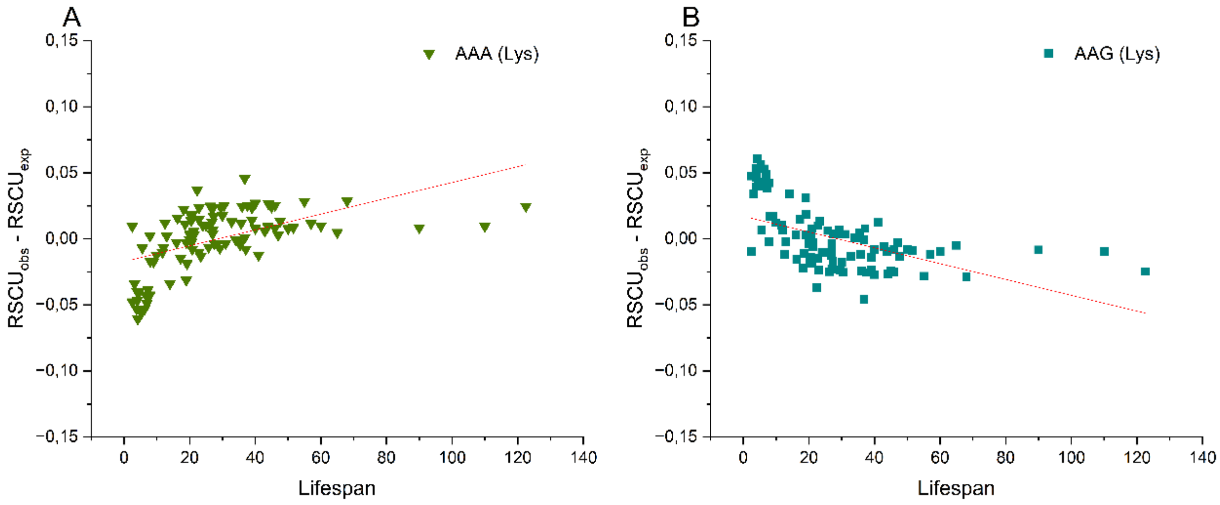


**Supplementary figure 12.b.** Differences between observed and expected relative synonymous codon usage (RSCU) values for codons encoding **lysine**, plotted against maximum lifespan (years) of 96 **mammalian** species.

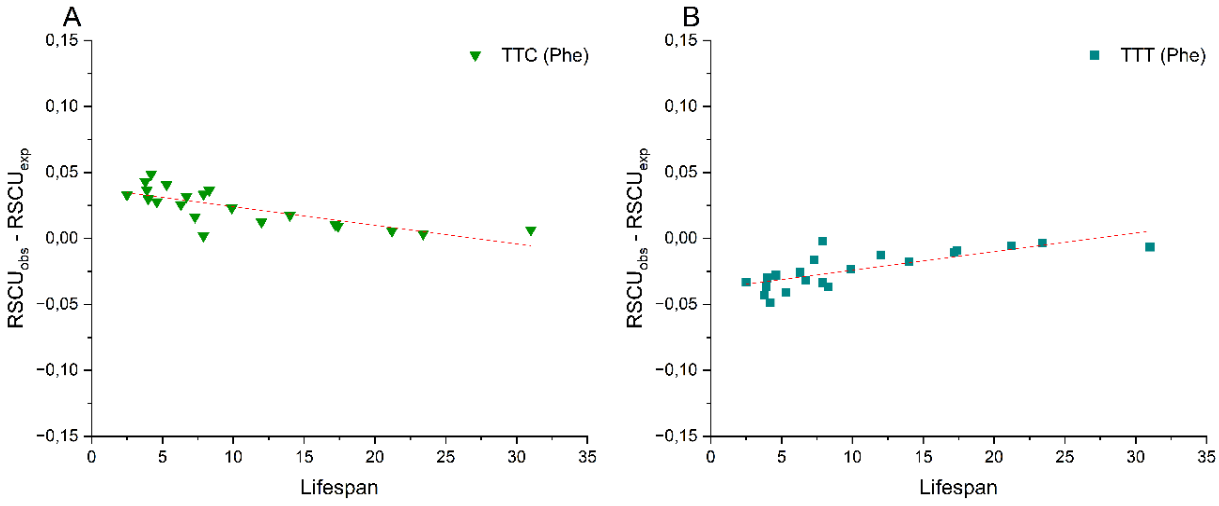


**Supplementary figure 13.a.** Differences between observed and expected relative synonymous codon usage (RSCU) values for codons encoding **phenylalanine**, plotted against maximum lifespan (years) of 19 **rodent** species.

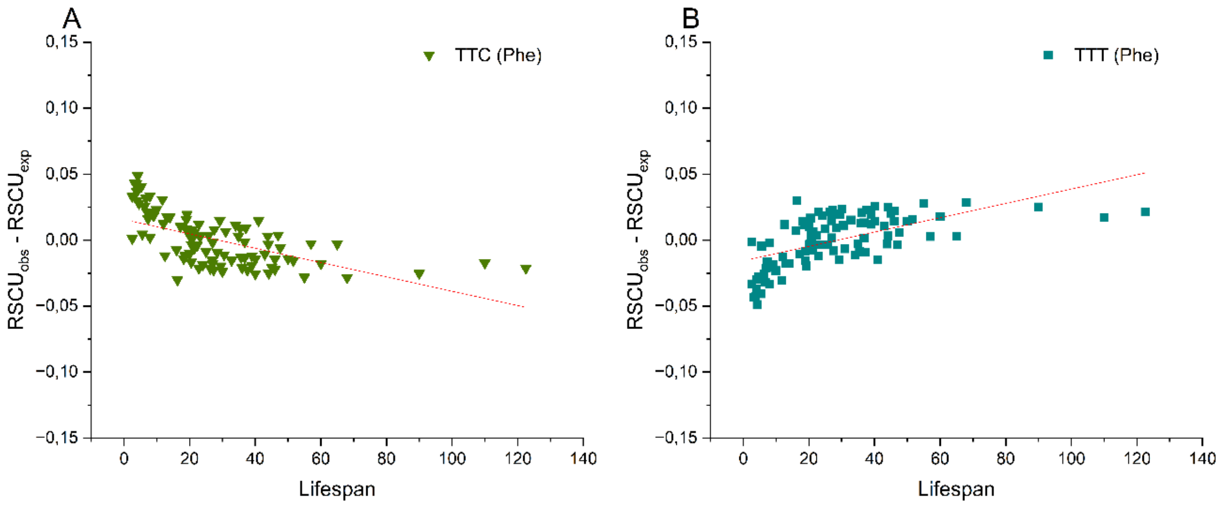


**Supplementary figure 13.b.** Differences between observed and expected relative synonymous codon usage (RSCU) values for codons encoding **phenylalanine**, plotted against maximum lifespan (years) of 96 **mammalian** species.

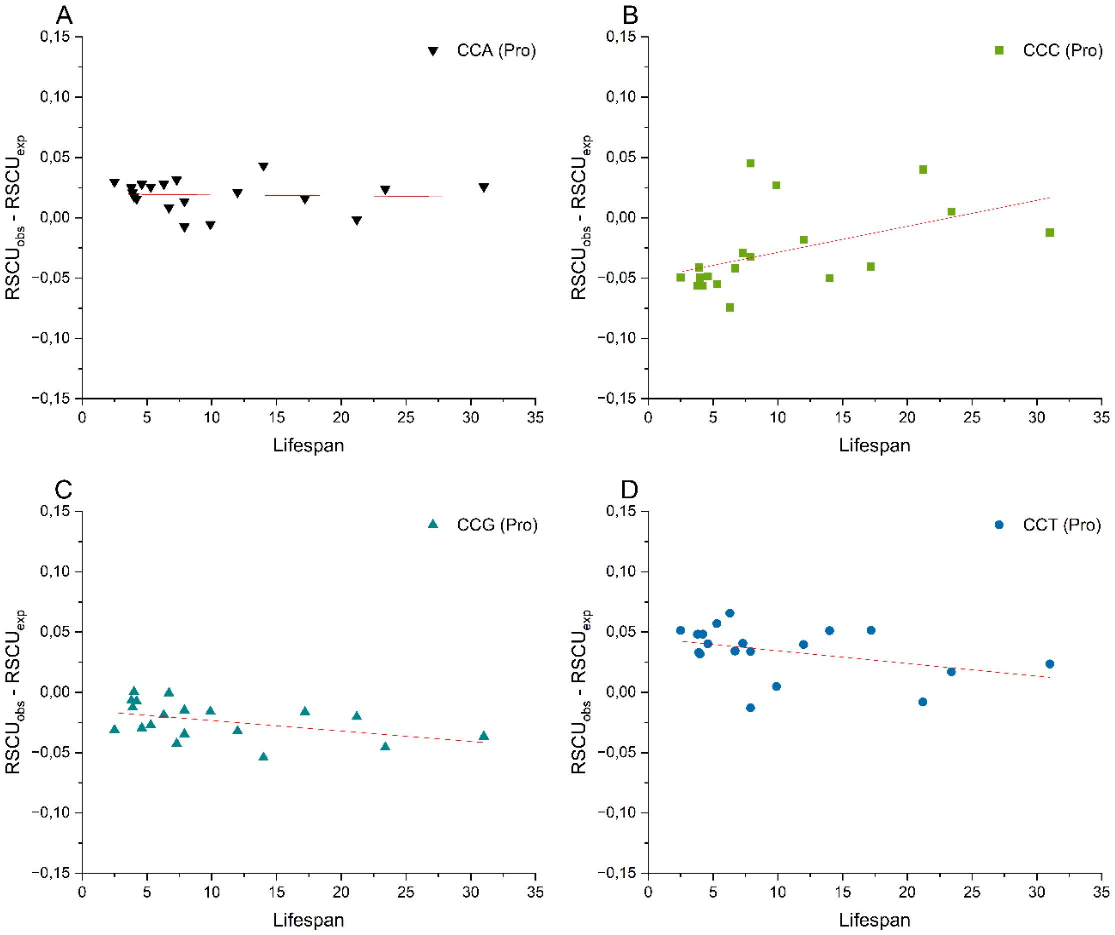


**Supplementary figure 14.a.** Differences between observed and expected relative synonymous codon usage (RSCU) values for codons encoding **proline**, plotted against maximum lifespan (years) of 19 **rodent** species.

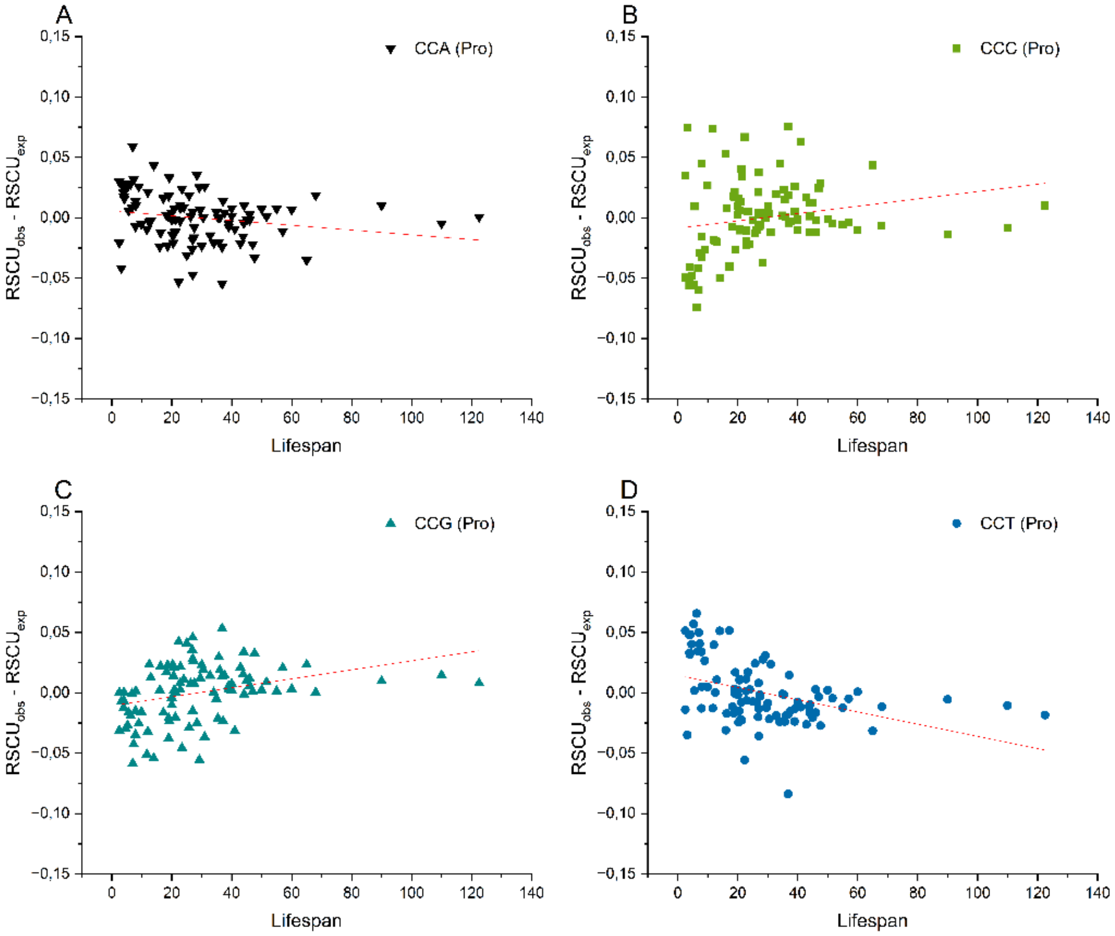


**Supplementary figure 14.b.** Differences between observed and expected relative synonymous codon usage (RSCU) values for codons encoding **proline**, plotted against maximum lifespan (years) of 96 **mammalian** species.

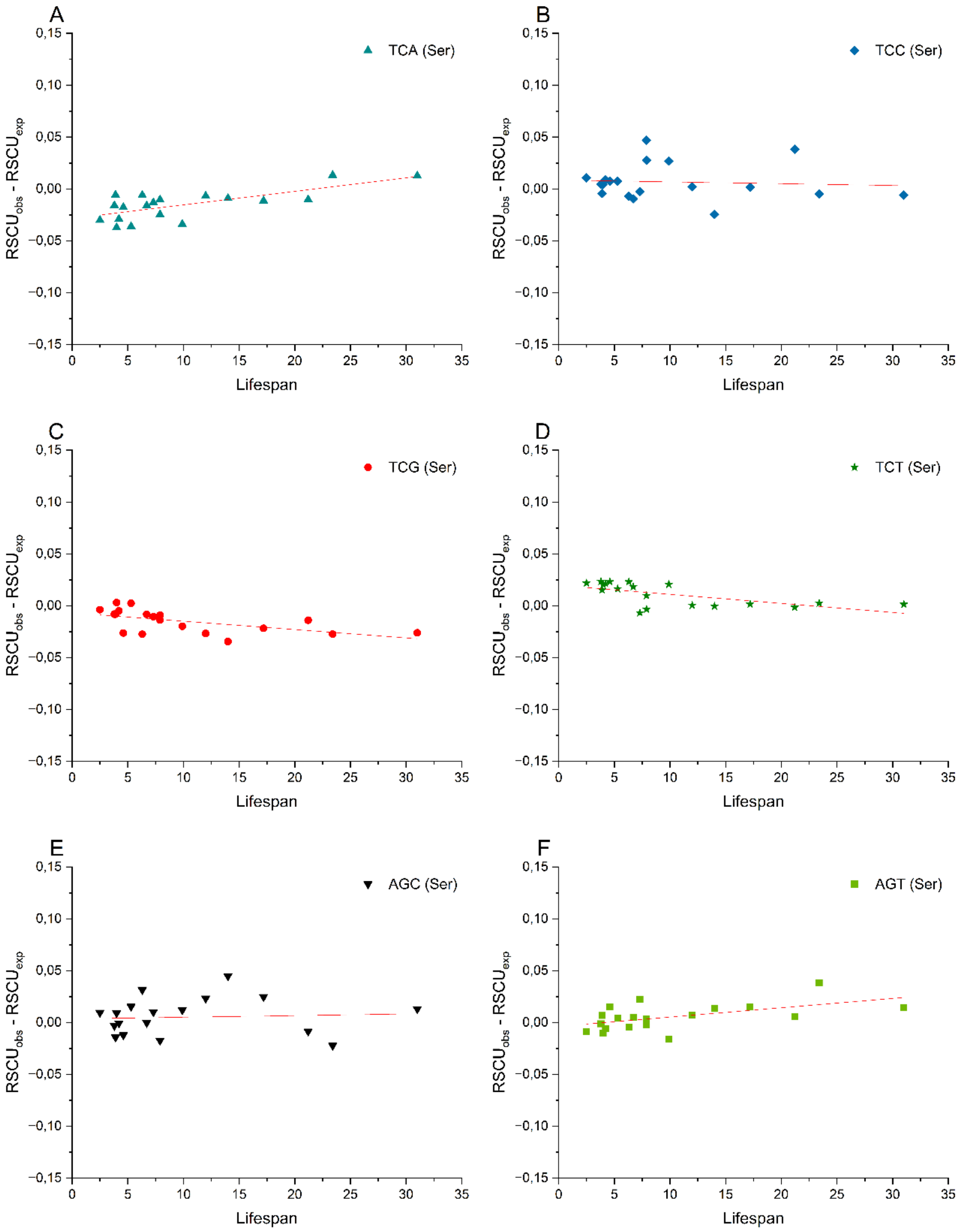


**Supplementary figure 15.a.** Differences between observed and expected relative synonymous codon usage (RSCU) values for codons encoding **serine**, plotted against maximum lifespan (years) of 19 **rodent** species.

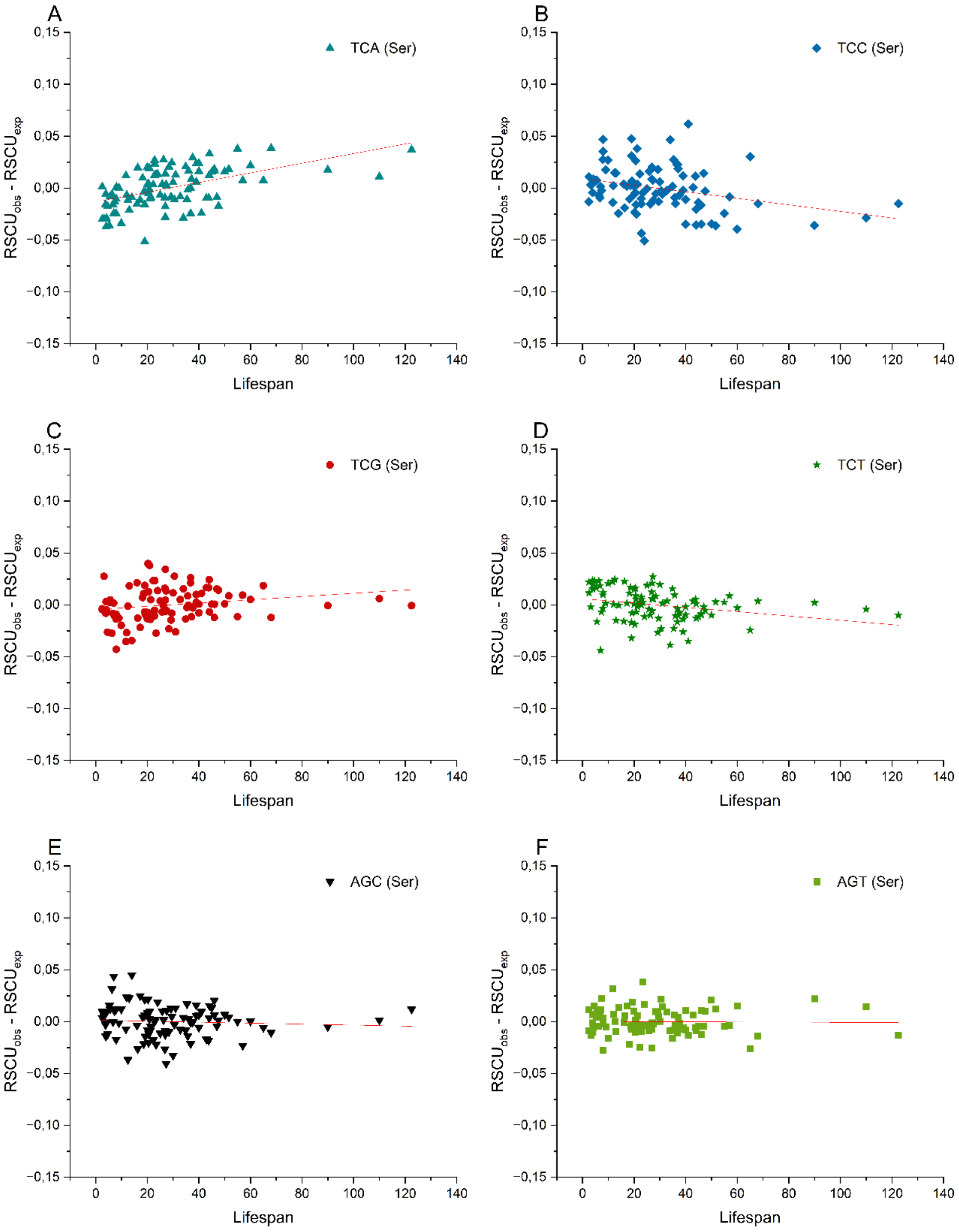


**Supplementary figure 15.b.** Differences between observed and expected relative synonymous codon usage (RSCU) values for codons encoding **serine**, plotted against maximum lifespan (years) of 96 **mammalian** species.

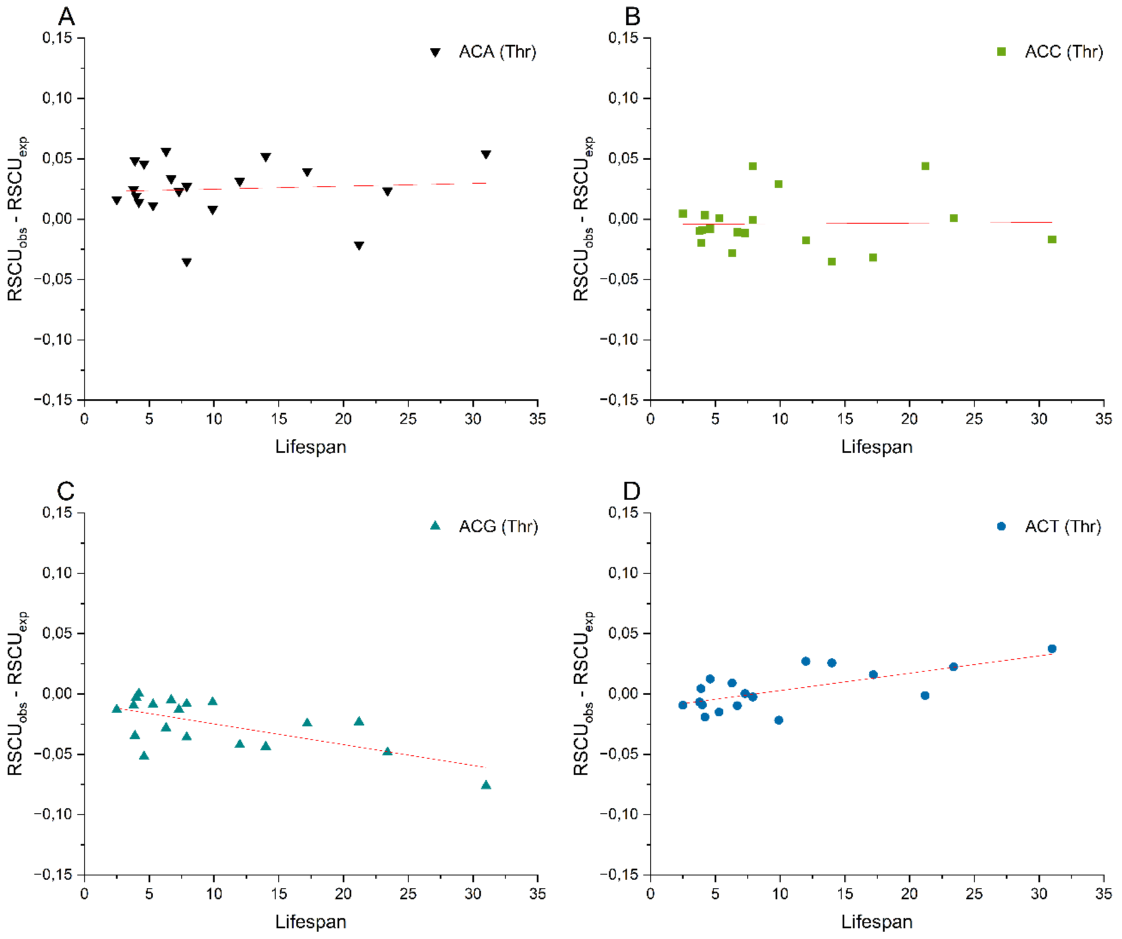


**Supplementary figure 16.a.** Differences between observed and expected relative synonymous codon usage (RSCU) values for codons encoding **threonine**, plotted against maximum lifespan (years) of 19 **rodent** species.

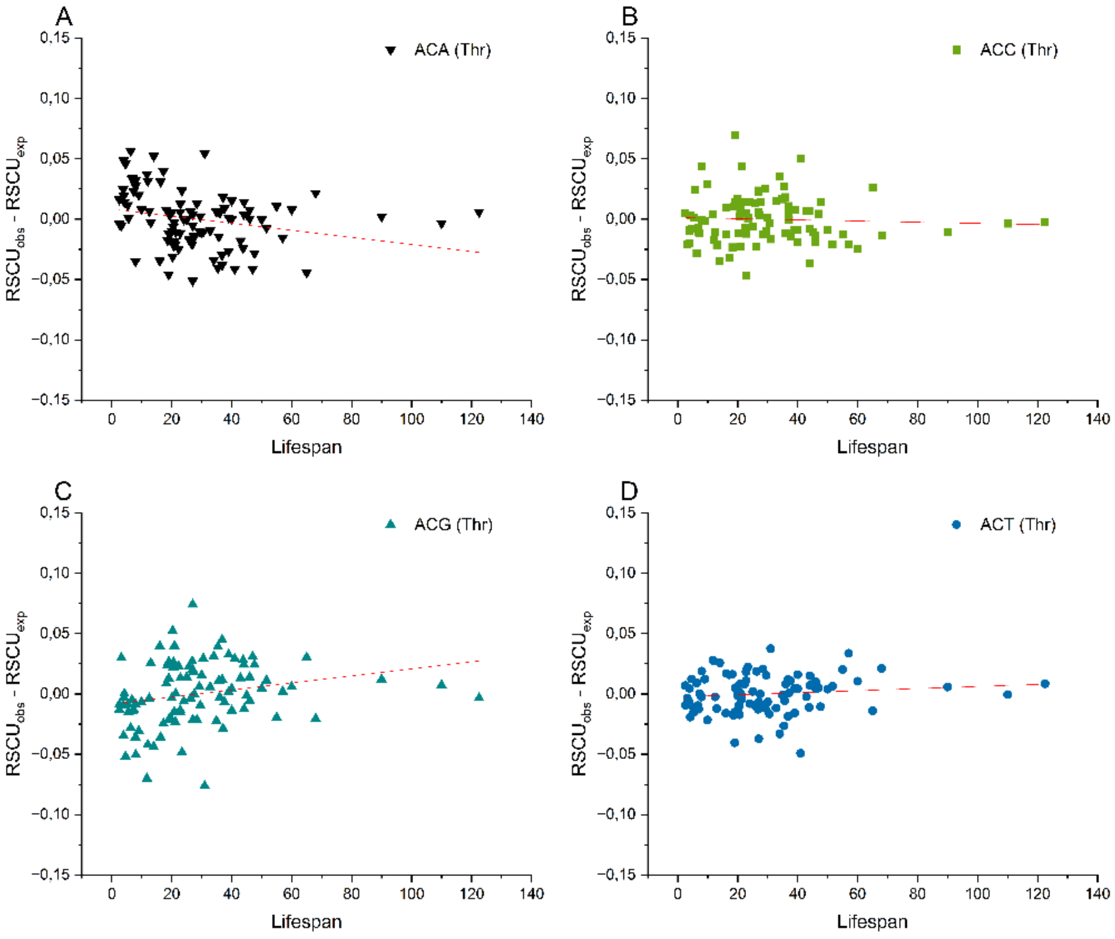


**Supplementary figure 16.b.** Differences between observed and expected relative synonymous codon usage (RSCU) values for codons encoding **threonine**, plotted against maximum lifespan (years) of 96 **mammalian** species.

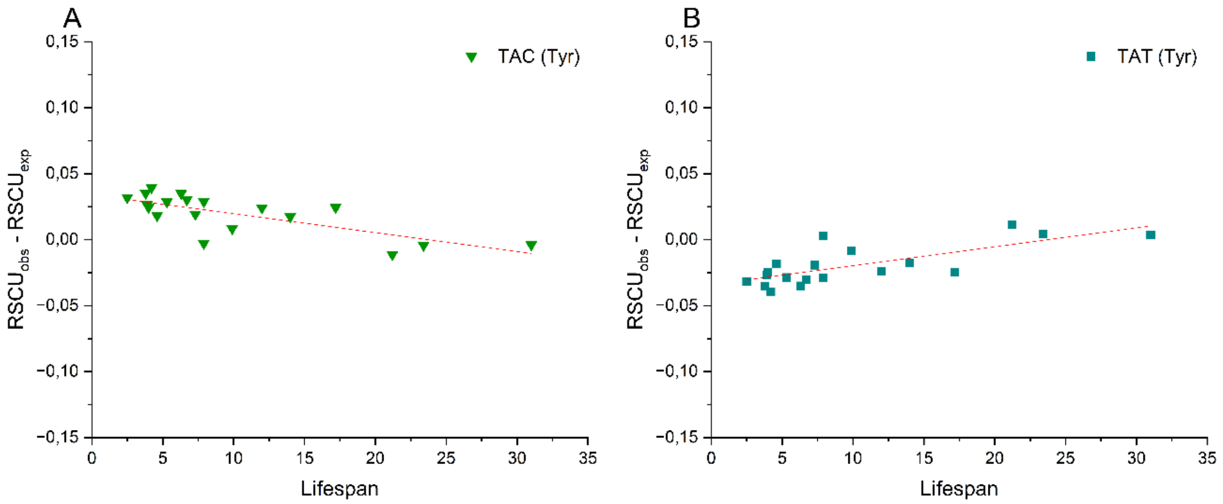


**Supplementary figure 17.a.** Differences between observed and expected relative synonymous codon usage (RSCU) values for codons encoding **tyrosine**, plotted against maximum lifespan (years) of 19 **rodent** species.

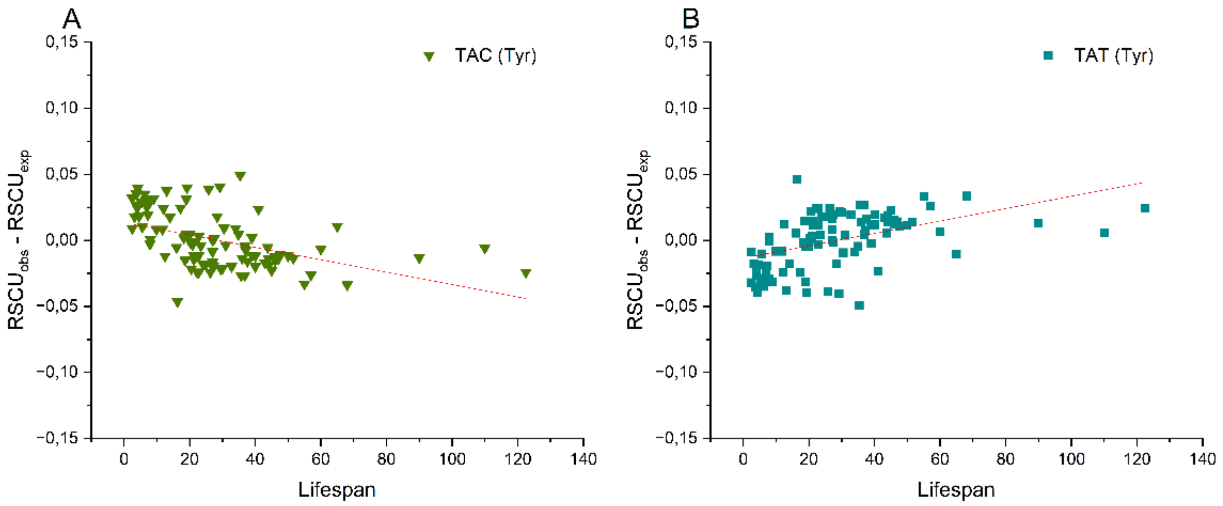


**Supplementary figure 17.b.** Differences between observed and expected relative synonymous codon usage (RSCU) values for codons encoding **tyrosine**, plotted against maximum lifespan (years) of 96 **mammalian** species.

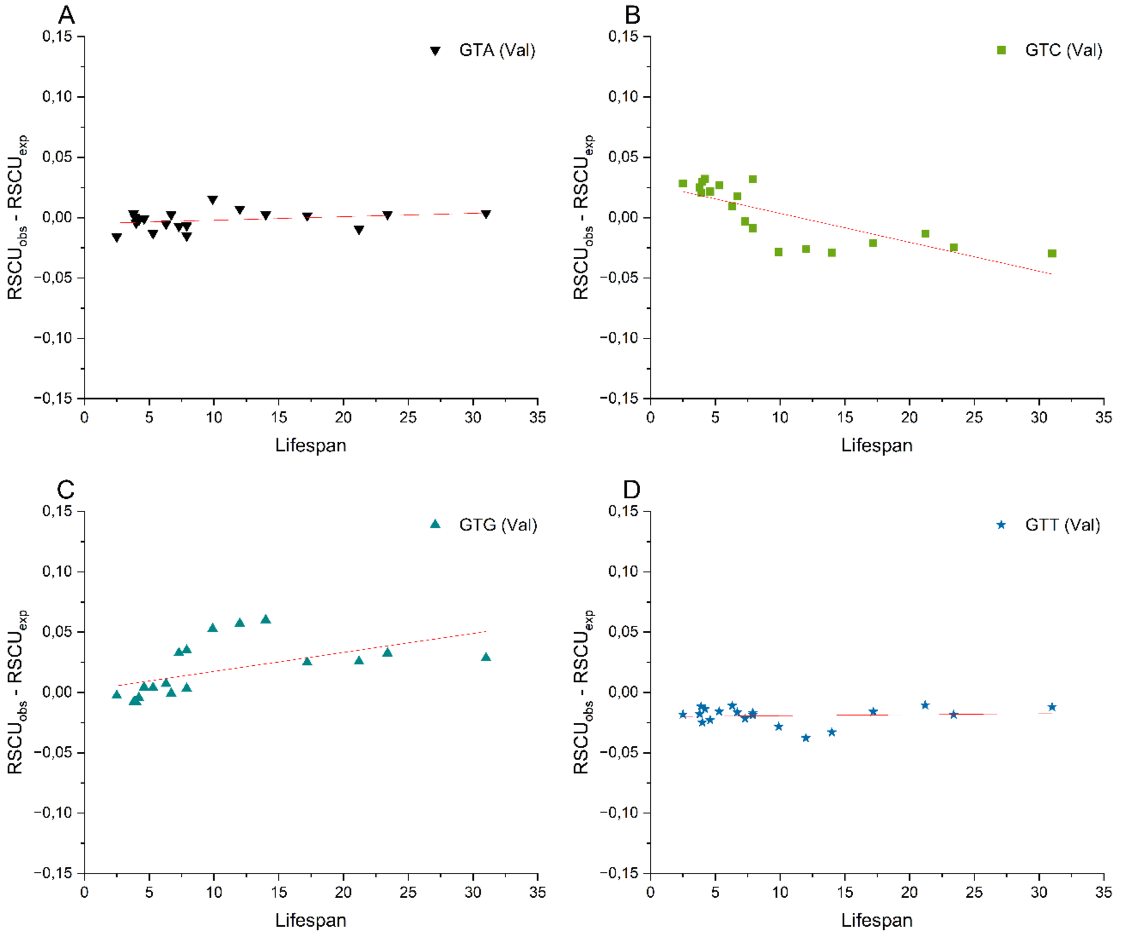


**Supplementary figure 18.a.** Differences between observed and expected relative synonymous codon usage (RSCU) values for codons encoding **valine**, plotted against maximum lifespan (years) of 19 **rodent** species.

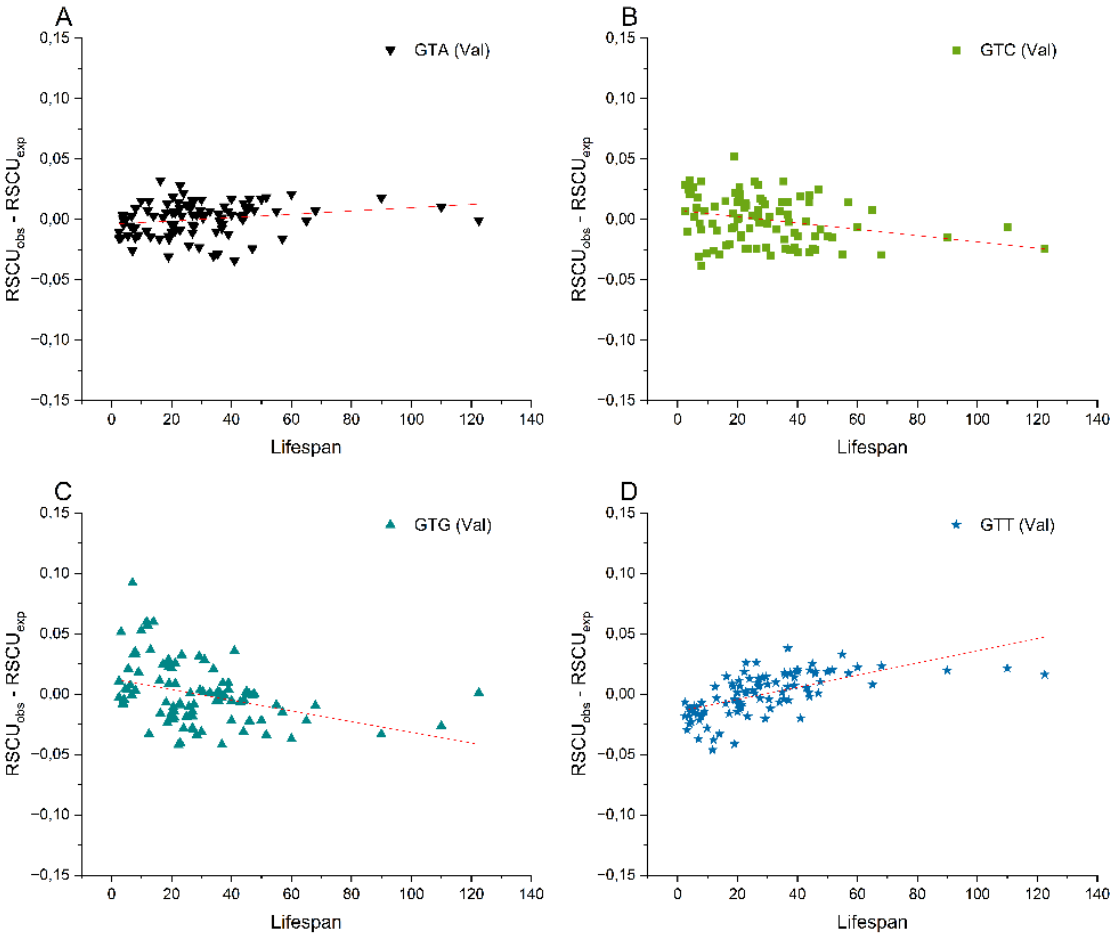


**Supplementary figure 18.b.** Differences between observed and expected relative synonymous codon usage (RSCU) values for codons encoding **valine**, plotted against maximum lifespan (years) of 96 **mammalian** species.

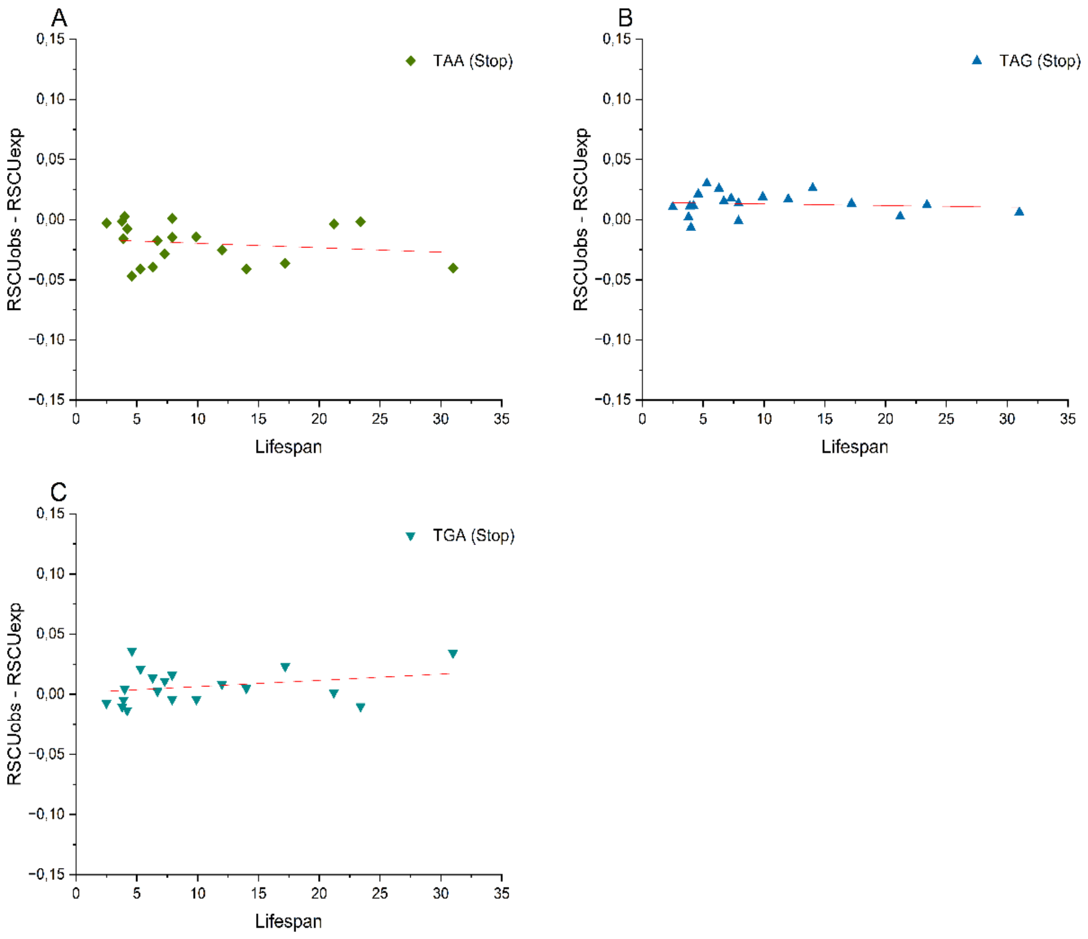


**Supplementary figure 19.a.** Differences between observed and expected relative synonymous codon usage (RSCU) values for **stop** codons, plotted against maximum lifespan (years) of 19 **rodent** species.

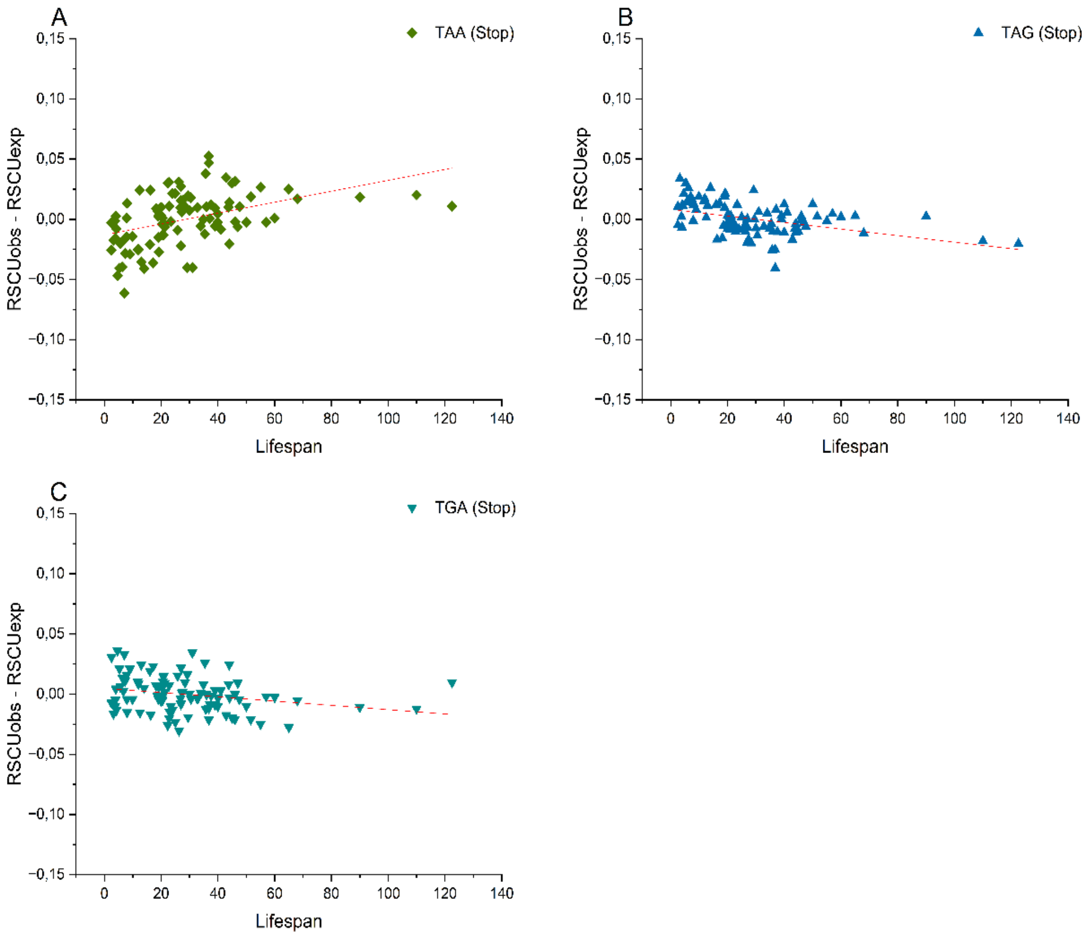


**Supplementary figure 19.b.** Differences between observed and expected relative synonymous codon usage (RSCU) values for **stop** codons, plotted against maximum lifespan (years) of 96 **mammalian** species.
